# Characterization of conformational dynamics and allostery of the catalytic domain of human mitochondrial YME1L protease

**DOI:** 10.1101/2025.01.30.635686

**Authors:** Megan K. Black, Angelina S. Kim, Ching Y. Chen, Monica M. Goncalves, Simrah Waseem, Siavash Vahidi, Rui Huang

## Abstract

Mitochondrial proteostasis is essential to maintain cellular function and survival. YME1L protease is an important contributor to proteostasis which belongs to the AAA+ (<u>A</u>TPases <u>A</u>ssociated with diverse cellular <u>A</u>ctivities) protease family and is anchored to the inner mitochondrial membrane. YME1L plays a pivotal role in mitochondrial protein quality control by selectively degrading misfolded and native proteins. The precise mechanisms by which nucleotide binding and hydrolysis influence YME1L’s conformational dynamics, proteolytic activity, and stability remain unclear. Here we characterize the conformational dynamics and allosteric regulation of the YME1L catalytic domain. Using a hexameric soluble YME1L construct, we employ hydrogen/deuterium exchange mass spectrometry (HDX-MS) and nuclear magnetic resonance (NMR) spectroscopy to demonstrate that nucleotide binding reduces the backbone flexibility and modulates the side-chain dynamics of the AAA+ domain, while Zn²⁺ binding stabilizes the protease domain. We also reveal novel long-range allostery between the AAA+ and protease domains of YME1L, mediated by a critical salt bridge on the inter-domain interface. We show the importance of the salt bridge in facilitating ATP-dependent substrate degradation by YME1L. Additionally, we show that ATP binding stabilizes the structure of the catalytic domain of YME1L and protects it from chemical modification- and heat-induced aggregation. These findings explain the nucleotide-driven regulation of YME1L and provide novel insights into understanding its proteolytic activity and structural stability under physiological and stress conditions.

## Introduction

Mitochondria contain approximately 1,100 unique proteins, with all but 13 encoded by nuclear DNA and translated in the cytoplasm (1, 2). These proteins are imported into the mitochondria, processed, and folded into their functional forms, while their quality is maintained by a complex network of pathways that regulate biosynthesis, import, folding, degradation, and mitophagy (3). This quality control system is essential for mitochondrial functionality and overall cell survival, as improper protein folding can lead to toxic aggregation. A key aspect of this quality control involves degradation of misfolded mitochondrial proteins, a process mediated by a family of AAA+ (<u>A</u>TPases <u>A</u>ssociated with diverse cellular <u>A</u>ctivities) proteases found in all mitochondrial compartments (4, 5). These proteases use the energy from ATP hydrolysis to unfold and degrade protein substrates.

One such mitochondrial protease, YME1L, is a membrane-anchored AAA+ protease that localizes to the inner mitochondrial membrane (IMM) with its catalytic domain extending into the intermembrane space (IMS). YME1L is a hexameric complex, with each subunit containing an N-terminal domain, a single transmembrane helix, followed by a catalytic domain that is soluble in absence of the transmembrane domain and consists of a AAA+ (<u>A</u>TPases <u>A</u>ssociated with diverse cellular <u>A</u>ctivities) domain and a Zn^2+^-dependent metalloprotease domain (Fig. 1A and B) (6, 7).

**Figure 1.**
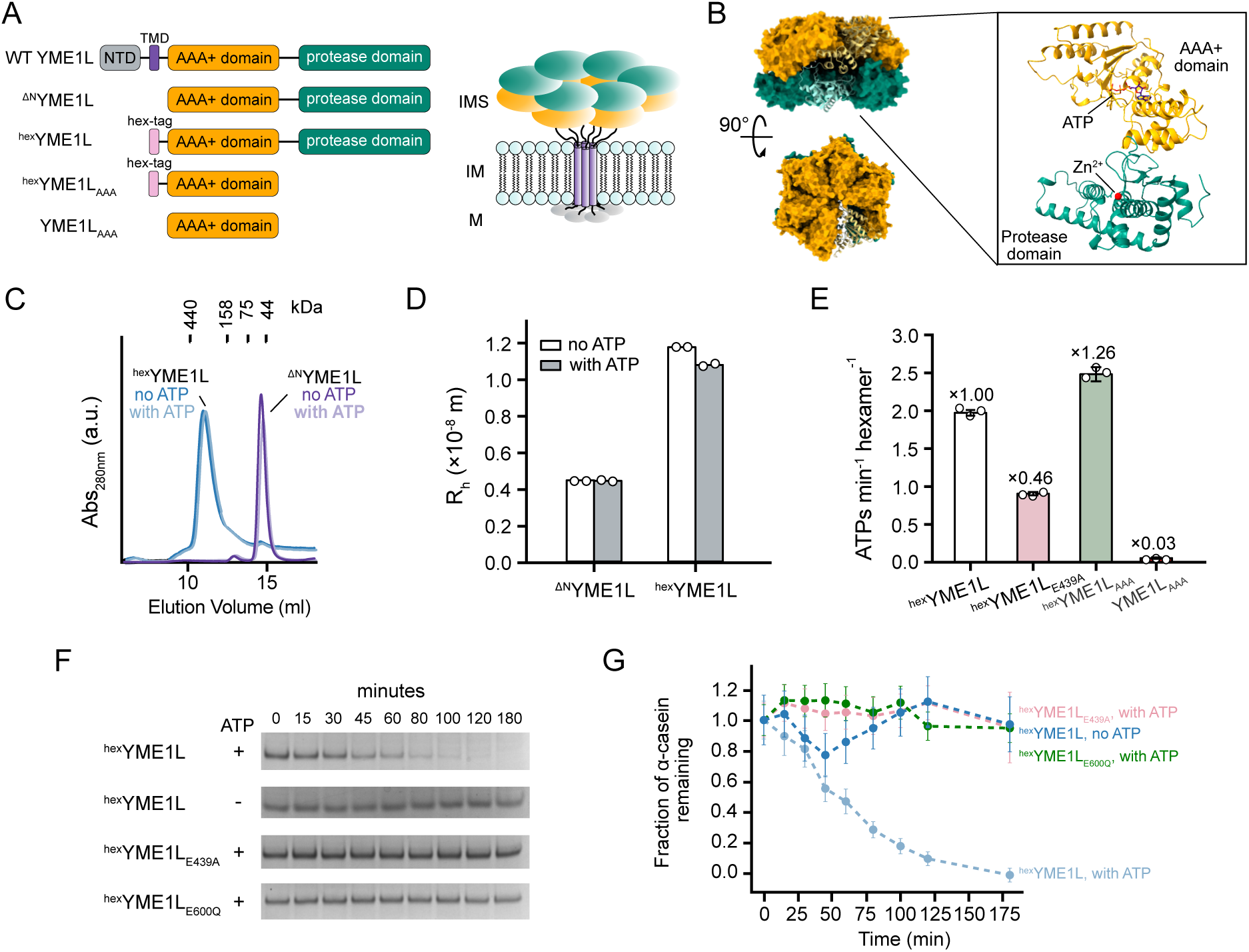
Oligomerization and enzymatic activity of the catalytical domain of YME1L in the presence of a hex-tag. **(A)** Left: Domain organization of a WT YME1L subunit (WT YME1L) and cartoon representations of the YME1L constructs used in the study. Right: Schematic representation of the hexameric YME1L anchored to the inner membrane of mitochondria (M = mitochondrial matrix; IM: mitochondrial inner membrane; IMS: mitochondrial intermembrane space). **(B)** Structural model of the hexameric catalytic domain (residue 317-773) of YME1L, predicted by AlphaFold3 (69), featuring two distinct layers – the AAA+ domains (dark yellow) and the protease domains (dark green), together enclosing the central pore. The predicted structure of a monomer bound to ATP and Zn^2+^ is enlarged. **(C)** SEC elution profiles of ^hex^YME1L and ^ΔN^YME1L in the presence and absence of 5 mM ATP. The molecular weights of the standards are given along the top. **(D)** Hydrodynamic radii of ^hex^YME1L and ^ΔN^YME1L in the presence and absence of 10 mM ATP obtained by dynamic light scattering. Data were measured at the subunit concentration of 200 μM, 15 °C. **(E)** Rate of ATP hydrolysis by ^hex^YME1L, ^hex^YME1L_E439A_ (Walker-B mutant), ^hex^YME1L_AAA_, and the monomeric YME1L_AAA_. Normalized ATPase activities with respect to that of ^hex^YME1L are denoted above the bars**. (F)** Degradation of α-casein by ^hex^YME1L, ^hex^YME1L_E439A_, and ^hex^YME1L_E600Q_ (protease-inhibiting mutant). **(G)** The fraction of remaining α-casein over time in the degradation assays shown in (F). Images of the uncropped SDS-PAGE gels are shown in Fig. S2.

The AAA+ domain of YME1L harnesses the chemical energy from ATP hydrolysis to drive unfolding and translocation of substrates through the central pore of the enzyme and into the protease domain for degradation (8). YME1L plays a vital role in maintaining mitochondrial proteostasis. Disruption of YME1L function leads to mitochondrial dysfunction and reduced cell survival under stress conditions (9, 10). YME1L is upregulated in several cancers, making it a promising target for cancer diagnosis and therapeutics (11–14). YME1L removes aberrant proteins from the IMM and IMS, such as the unassembled subunits of respiratory Complex I (*e.g.*, NDUFB6 and ND1) and Complex IV (*e.g*., COX4) (4, 6, 15), and degrades specific native mitochondrial proteins to adapt to cellular stress. Under stress conditions YME1L degrades TIMM17A, a component of the protein translocase TIM23 complex in the IM, in order to reduce protein import (16). Additionally, YME1L regulates mitochondrial morphology by processing OPA1, a dynamin-like protein, which exists in both a long membrane-bound form (L-OPA1) and a soluble short form (S-OPA1) in the mitochondria (17). The balance between these two forms is crucial for maintaining mitochondrial morphology (18–20). YME1L and the ATP-independent protease OMA1 cleave L-OPA1 to produce S-OPA1, particularly under stress conditions such as membrane depolarization and oxidative stress, promoting mitochondrial fission (21–23). On the other hand, high ATP levels activate YME1L to degrade OMA1, counteracting OMA1-induced mitochondrial fragmentation, while depletion of ATP leads to inactivation of YME1L, allowing it to be reciprocally degraded by OMA1 (9). It was suggested that conformational changes driven by ATP binding protect YME1L from hydrogen peroxide-induced modification (24) and prevent YME1L from degradation by OMA1 under oxidative stress (9). However, the precise mechanisms by which nucleotide binding and hydrolysis affect YME1L’s conformational dynamics and stability are not fully understood.

Here we present a detailed characterization of the conformational dynamics, allosteric modulation, and stability of the catalytic domain of YME1L. Using size-exclusion chromatography (SEC) and dynamic light scattering (DLS) we demonstrate that isolated soluble domain of YME1L has a low propensity to oligomerize. Using a hexameric soluble construct of YME1L, we characterize the nucleotide- and Zn^2+^-induced changes in conformational dynamics using a combination of hydrogen/deuterium exchange mass spectrometry (HDX-MS) and nuclear magnetic resonance (NMR) spectroscopy. Our results show that ATP and ADP binding significantly reduce backbone flexibility in the AAA+ domain, while Zn^2+^ binding leads to a rigidification of the protease domain. ADP binding also modulates the side-chain dynamics of the AAA+ domain. Furthermore, we uncover long-range functional allosteric communication between the AAA+ and protease domains. Finally, we demonstrate that ATP binding enhances YME1L’s thermal stability and protects it from aggregation due to chemical modification. Our findings offer insights into how nucleotide and cofactor binding modulate YME1L’s conformational dynamics and stability in vitro.

## Results

### A hexamerization tag promotes the formation of active hexameric YME1L catalytic domain

The ATPase and ATP-dependent protease activity of YME1L require its hexameric form. YME1L oligomerization, in turn, is facilitated by its N-terminal domain (NTD) and transmembrane domain (TMD) (Fig. 1A) (6). Previous studies showed that replacing the NTD and TMD with a soluble 32-residue hexameric coiled-coil domain promotes the hexamerization of the YME1L catalytic domain (^ΔN^YME1L) and restore its ATPase and protease activities (6, 8). Following this approach, we used the same construct consisting of an N-terminal hex-tag connected to the catalytic domain of YME1L by a 10-residue linker (^hex^YME1L) (Fig. 1A) and characterized its hydrodynamic properties and enzymatic activity.

We first used size exclusion chromatography (SEC) to characterize the oligomeric states of ^hex^YME1L and ^ΔN^YME1L. The SEC peak of ^hex^YME1L is centred at 11 mL while that of ^ΔN^YME1L is at 15 mL (Fig. 1C). These elution volumes are consistent with the molecular weights of the hexameric (332 kDa) and monomeric YME1L (51 kDa), respectively, consistent with previous findings (6). The elution volume of ^ΔN^YME1L remained unchanged (*i.e.*, monomeric) in the presence of 5 mM ATP (Fig. 1C), suggesting that addition of nucleotide alone is insufficient to promote hexamerization of the catalytic domain in the absence of the hex-tag at low protein concentrations (∼6 μM monomeric concentration at the center of the SEC peak). We subsequently used dynamic light scattering (DLS) to probe the concentration- and temperature-dependence of oligomerization for our YME1L constructs. We measured the hydrodynamic radius of ^ΔN^YME1L and ^hex^YME1L in the presence and absence of 10 mM ATP with the monomeric concentration of YME1L ranging from 25 to 200 μM, recorded at temperatures from 7.5 to 50 °C. We observed no changes in hydrodynamic radius between the nucleotide-free and ATP-bound states of ^ΔN^YME1L at the tested concentration and temperature ranges prior to onset of protein aggregation (Fig. 1D and Fig. S1). This suggests that the catalytic domain of YME1L has low intrinsic propensity to oligomerize even in the presence of nucleotide. This observation highlights the crucial role that the N-terminal and transmembrane domains play in the proper assembly of the YME1L hexamer.

We measured the ATPase activity of our constructs using an NADH-coupled assay (25). The initial ATPase rate for ^hex^YME1L was 1.97 ± 0.04 min⁻¹ · hexamer⁻¹ (Fig. 1E), consistent with a previous report (6). In contrast, mutating the Walker-B motif E439 to an alanine (^hex^YME1L_E439A_) leads to a 46% reduction in the ATPase activity compared to that of ^hex^YME1L (Fig. 1E), in agreement with a conserved mechanism for ATP hydrolysis in AAA+ motors. In addition, the hex-tag together with the AAA+ domain of YME1L (^hex^YME1L_AAA_) displays ∼26% higher ATPase activity compared with ^hex^YME1L (Fig. 1E), suggesting that the AAA+ domain itself is sufficient to form the obligate hexamer to support ATP hydrolysis in the presence of the hex-tag. The AAA+ domain without the hex-tag (YME1L_AAA_) showed no measurable ATPase activity, likely due to the monomeric state of the AAA+ domain (Fig. 1E). We also characterized the protease activity of ^hex^YME1L using α-casein as a model substrate. ^hex^YME1L degrades α-casein in an ATP-dependent manner (Fig. 1F and G, and Fig. S2). In contrast, a YME1L mutant with a catalytically deficient protease active site (^hex^YME1L_E600Q_), in which the glutamate in the HE*XX*H Zn^2+^-binding motif is mutated to glutamine (26, 27), showed no protease activity in the presence of ATP (Fig. 1F and G, and Fig. S2). Furthermore, we observed no appreciable degradation of α-casein by the Walker B mutant (^hex^YME1L_E439A_) (Fig. 1F and G, and Fig. S2). These results demonstrate that the hex-tag effectively mimics the hexamerization function of NTD and TMD, thereby enabling the ^hex^YME1L construct to perform ATP-dependent substrate translocation and degradation.

### Nucleotide binds to monomeric AAA+ domain of YME1L and modifies side chain dynamics of the AAA+ domain

We employed solution NMR spectroscopy to examine the effect of nucleotide binding on the side-chain dynamics of YME1L. Given the large molecular weight of YME1L (51 kDa for the monomeric ^ΔN^YME1L and 330 kDa for ^hex^YME1L), we isotopically labeled the methyl groups of Ile (δ1), Leu (δ1,δ2), Val (γ1,γ2), and Met (ε) side chains to introduce NMR-detectable ^13^CH_3_-methyls in an otherwise highly deuterated background. This labeling scheme, referred to in what follows as [^13^CH_3_-ILVM], in combination with NMR experiments that exploit the methyl-TROSY effect (28–30) allowed us to achieve great sensitivity and resolution in the ^1^H-^13^C correlated NMR spectra of the monomeric YME1L_AAA_ (Fig. 2B and Fig. S3B) and the monomeric ^ΔN^YME1L (Fig. 2F, and Fig. S3C). However, [^13^CH_3_-ILVM] labeled ^hex^YME1L_AAA_ failed to yield high-quality NMR spectra, potentially due to dynamic oligomerization and structural heterogeneity among subunits (Fig. S4A). The spectral quality improves upon the addition of an ATP analogue, ATPγS (Fig. S4B), yet the resolution is not sufficient for further investigation. Therefore, our NMR characterization focuses on the monomeric YME1L constructs. Although we could not obtain the identity of each methyl peak due to the low yield of the isotopically labeled YME1L, we were able to characterize the binding of ADP and its effect on the dynamics of the methyl side chains in our monomeric YME1L constructs. We first titrated ADP into [^13^CH_3_-ILVM] labeled YME1L_AAA_, which resulted in the appearance of two sets of peaks, corresponding to the ligand-free and ligand-bound populations (Fig. S5A). This observation indicates that ADP binds to the monomeric AAA domain and that the binding causes the so-called “slow chemical exchange” in the NMR spectra (31, 32). This phenomenon can be exploited to quantify concentrations of the free and bound forms of protein. By fitting the titration profiles of the free and bound peaks, we determined an average dissociation constant *K_d_* value of 4.6 ± 2.3 μM (Fig. 2B and Fig. S3D), consistent with previously reported values (33). Interestingly, most residues displayed an increase in peak intensities in the ADP-bound state compared to the apo state (Fig. 2D), suggesting reduced structural inhomogeneity or decreased millisecond timescale dynamics in the AAA+ domain upon ADP binding. We subsequently recorded the ^1^H-^13^C correlated spectra of [^13^CH_3_-ILVM] labeled ^ΔN^YME1L. The spectrum of YME1L_AAA_ largely overlaps with that of ^ΔN^YME1L (Fig. 2E and Fig. S6), allowing us to distinguish peaks that originate from the AAA+ domain (present in both YME1L_AAA_ and ^ΔN^YME1L spectra) from those that originate from the protease domain (only present in the ^ΔN^YME1L spectra). Addition of ADP to the monomeric ^ΔN^YME1L induced chemical shift changes in peaks originating from the AAA+ domain (Fig. 2F), similar to those observed in the monomeric YME1L_AAA_ (Fig. 2B). Additionally, intensity increase was primarily observed in the AAA+ domain with minimal changes in the protease domain (Fig. 2G), indicating that nucleotide binding primarily alters side-chain dynamics within the AAA+ domain.

**Figure 2.**
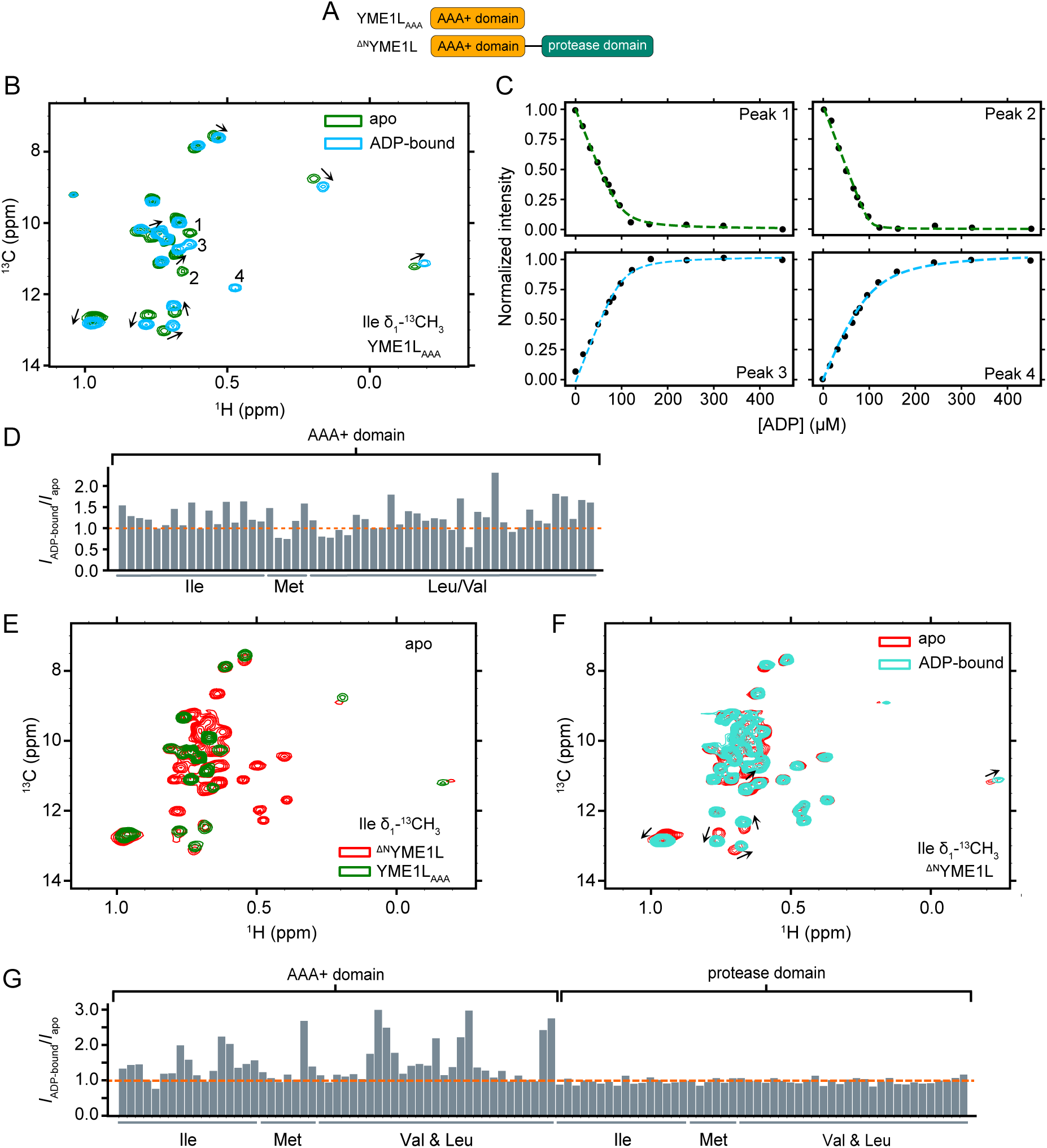
Nucleotide binding alters side chain dynamics in the AAA+ domain. **(A)** Domain organization of the two constructs, YME1L_AAA_ and ^ΔN^YME1L, presented in the figure. **(B)** Overlay of ^13^C-^1^H HMQC spectra of [U-^2^H, ILVM-^13^CH_3_] labeled YME1L_AAA_ in the presence (blue) and absence (green) of 5 mM ADP. **(C)** Titration profiles of selected peaks from (B) with increasing concentration of ADP, fitted to a one-to-one binding model for slow chemical exchange. **(D)** Intensity ratios of the methyl peaks in YME1L_AAA_ in the ADP-bound versus apo states. The orange dashed line indicate an intensity ratio of 1.0. **(E)** Overlay of ^1^H-^13^C HMQC spectra of [U-^2^H, ILVM-^13^CH_3_] labeled YME1L_AAA_ (green) and ^ΔN^YME1L (red), respectively, in the apo state. **(F)** Overlay of ^13^C-^1^H HMQC spectra of [U-^2^H, ILVM-^13^CH_3_] labeled ^ΔN^YME1L in the presence (green) and absence (red) of 5 mM ADP. **(G)** Intensity ratios of the methyl peaks in ^ΔN^YME1L in the ADP-bound versus apo states. Spectra were acquired at 14.1 T and 20 °C.

### Backbone dynamics of YME1L is modulated by nucleotide and Zn^2+^ binding

We then used hydrogen/deuterium exchange mass spectrometry (HDX-MS) to examine changes in backbone dynamics across various cofactor-bound states. HDX-MS is highly sensitive to molecular motions, as the HDX rate of individual backbone amides is influenced by their solvent exposure and participation in hydrogen bonding networks (34). Rigid protein segments undergo HDX slowly, while regions that undergo local conformational fluctuations and transient opening of the hydrogen bonds become deuterated faster (34). HDX-MS is a well-established tool for investigating the conformational dynamics of AAA+ motor proteins through comparative analyses of cofactor-free and cofactor-bound states (35). Regions involved in ligand binding typically show a decrease in deuterium uptake, while allosteric changes in the protein can result in either an increase or decrease in uptake (36).

We performed bottom-up continuous labeling HDX experiment to monitor the structural transitions that hexameric YME1L undergoes in the apo, ADP-bound, ATP-bound, and ATP+Zn²⁺-bound forms. To reduce the rate of ATP hydrolysis during labeling, experiments were carried out on the ^hex^YME1L_E439A_ construct. Our peptide mapping strategy identified 157 high-quality peptides with quantifiable deuterium uptake levels, corresponding to 91% sequence coverage and a redundancy level of 4.6 (Fig. S7). The majority of these peptides displayed unimodal symmetric isotopic profiles that gradually incorporate deuterium over time. This is indicative of EX2 behavior where the HDX rate (*k*_HDX_) is significantly slower than local protein motions (*k*_cl_). We compared deuterium uptake along YME1L’s sequence across different cofactor-bound states relative to the apo form and presented the results as heat maps (Fig. 3). In both the ADP-bound (Fig. 2A) and ATP-bound (Fig. 3B) states, a series of peptides in the AAA+ domain showed reduced deuterium uptake, suggesting that nucleotide binding largely increases backbone rigidity in this domain. This is consistent with our NMR results where modulation of side chain dynamics was predominantly observed in the AAA+ domain in response to nucleotide binding. Peptides surrounding the nucleotide-binding pocket, in particular, those containing the Walker-A motif (peptides 377-387, 377-388, 378-387, 379-387, and 381-387) showed significant reduction in deuterium uptake in the ADP- and ATP-bound states, consistent with the function of the conserved K385 in the Walker-A motif coordinating the β-phosphate of nucleotides (37–39). Additionally, peptides within the small AAA+ sub-domain (residue 509-579) showed decreased deuterium uptake upon nucleotide binding, indicative of loss of conformational dynamics. Previous studies suggest that nucleotide binding, occurring in the cleft between the large and small AAA+ subdomains, induces rotations between the domains (39, 40), which may lead to the observed rigidification. The second region of homology (SRH) also exhibited reduced flexibility in nucleotide-bound states (peptides 468-488), consistent with its function of interacting with the nucleotide from a neighboring subunit (*in trans*) (41). Interestingly, a comparison between ADP- and ATP-bound states revealed that ADP-bound YME1L showed greater protection (reduced deuterium uptake) in the AAA+ domain, suggesting enhanced backbone rigidity upon nucleotide hydrolysis (Fig. 3C and Fig. S8).

**Figure 3.**
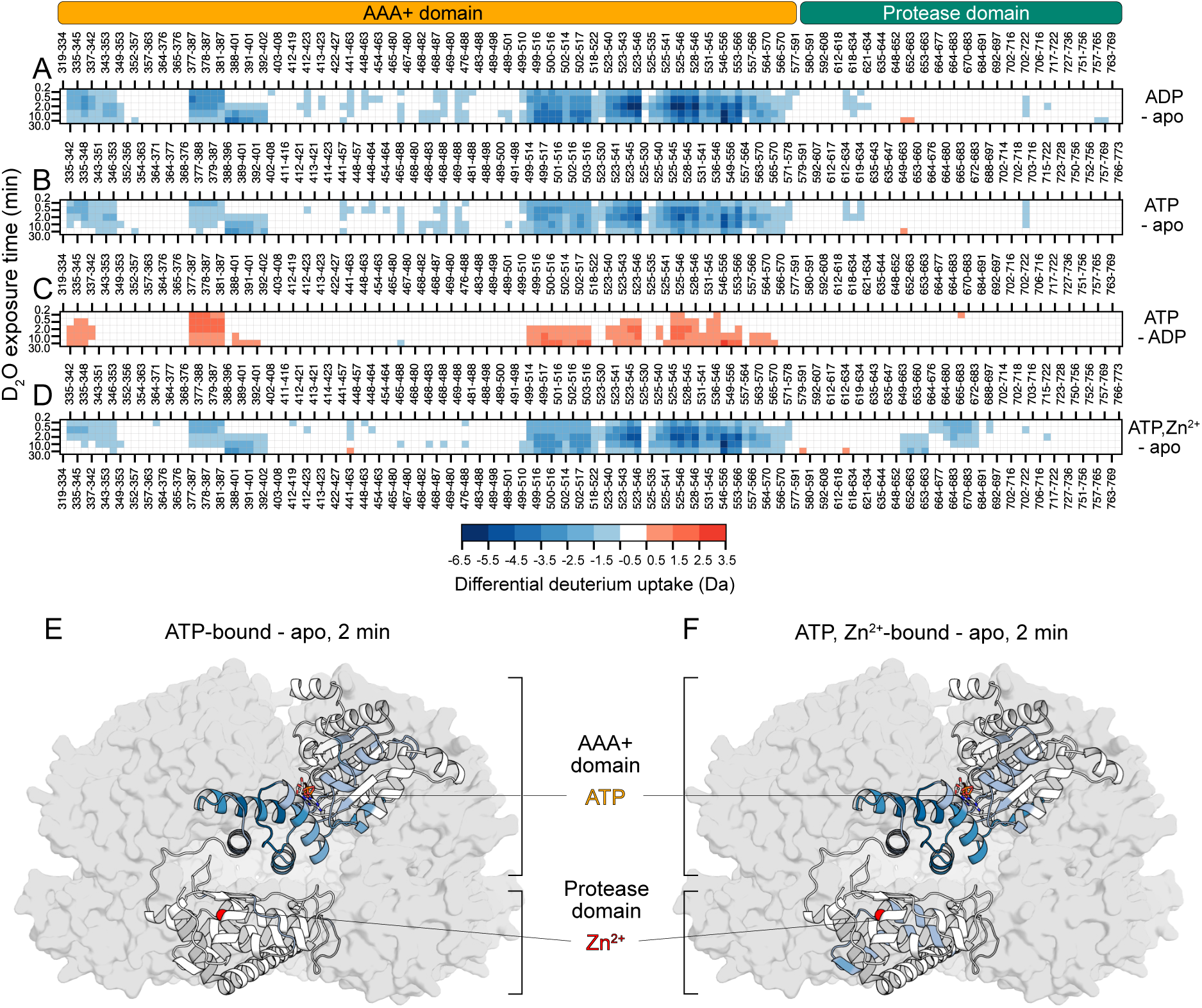
The structural dynamics of ^hex^YME1L are impacted by the binding of cofactors. Heatmaps display the differential deuterium uptake of ^hex^YME1L bound to **(A)** ADP and **(B)** ATP relative to apo ^hex^YME1L. Rigidification is observed across the AAA+ domain, mainly pronounced in regions that line the nucleotide binding site; **(C)** The deuterium uptake of the ATP-bound state compared to ADP-bound ^hex^YME1L suggests that the nucleotide binding site is more dynamic when bound to ATP and rigidifies upon hydrolysis; **(D)** The differential deuterium uptake of ATP and Zn^2+^-bound ^hex^YME1L relative to apo ^hex^YME1L shows a reduction of deuterium uptake in the nucleotide binding site of the AAA+ domain and the Zn^2+^ binding site of the protease domain; **(E)** The differential deuterium uptake of ATP-bound relative to apo ^hex^YME1L after 2 minutes of deuterium exposure are mapped onto an AlphaFold3 (69) structural model of the protein, emphasizing the reduction of deuterium uptake localized in the nucleotide binding site of the AAA+ domain; **(F)** Upon the addition of Zn^2+^, rigidification is observed of residues that line the Zn^2+^ binding site in the protease domain. Regions of the protein that are not covered during HDX-MS are shown in grey on cartoon representation of a monomer.

YME1L is a zinc metalloprotease, with active site residues H599, H603, and D677 coordinating a Zn²⁺. Unlike the ATPase domain, the protease domain exhibits little change in response to nucleotide binding. Interestingly, we observed a decrease in deuterium uptake for several peptides spanning residues 612-634 and 702-722 in the protease domain upon ADP or ATP binding (Fig. 3A and B). Residues 612-634 are located at the interface between the AAA+ and protease domains, while residues 702-722 are situated at the subunit interface and at the central pore (Fig. S9). This observation is consistent with allosteric communication between the AAA+ and protease domains. To probe the impact of hexamerization on this allosteric communication, we compared the deuterium uptake of the nucleotide-bound and apo states of a monomeric construct of YME1L (^ΔN^YME1L) (Fig. S10). Consistent with the HDX-MS results from the hexameric form (^hex^YME1L), nucleotide binding induces rigidification of the AAA+ domain of monomeric YME1L (^ΔN^YME1L). A comparison between ^hex^YME1L and ^ΔN^YME1L in the ATP-bound state revealed only slight differences, with ^hex^YME1L showing marginally more protection throughout the entire sequence (Fig. S10C), likely due to the hex-tag-promoted oligomerization (Fig. 1). Notably, we only detected the allosteric rigidification of the protease domain (peptides 612-634 and 702-722) in the ^hex^YME1L construct but not in the ^ΔN^YME1L construct, suggesting that hexamerization is required for allosteric communication between the AAA+ and protease domains of YME1L. Furthermore, comparing the deuterium uptake of the ATP, Zn²⁺-bound state of ^hex^YME1L to that of apo ^hex^YME1L also showed significantly reduced deuterium uptake in the Zn^2+^-binding region (residues 649-683), as anticipated (Fig. 3D).

### Functional allostery between the AAA+ and protease domains of YME1L

To explore functional allostery between the AAA+ and protease domains, we introduced an E600Q mutation at the protease active site of ^hex^YME1L to suppress proteolytic activity, and we tested the ATPase activity of the enzyme. Notably, this mutation resulted an 8.6-fold increase in the ATPase activity despite that the mutation site is located more than 40 Å from the ATP-binding site (Fig. 4A and B). This confirms long-range allosteric communication between the two domains. The presence or absence of Zn^2+^ in the protease active site does not significantly affect the ATPase activity (Fig. S11), consistent with the HDX-MS data showing that Zn^2+^ binding does not alter the deuterium uptake pattern of the AAA+ domain (Fig. 3B and D). To further investigate the allosteric effect of the E600Q mutation on the conformational dynamics of the AAA+ domain, we performed HDX-MS experiments on the E600Q mutant of ^hex^YME1L in both the ATP-bound and apo states (Fig. 4C and D). Upon ATP binding, we observed a reduction in deuterium uptake in the AAA+ domain (Fig. 4D), similar to that observed in ^hex^YME1L. However, compared to ^hex^YME1L, the differences in deuterium uptake between the ATP-bound and apo states were significantly smaller due to the E600Q substitution (Figs. 2B and 5C), suggesting the mutation at the proteolytic site alters the backbone dynamics of the AAA+ domain, potentially contributing to the 8.6-fold increase in the ATPase activity.

**Figure 4.**
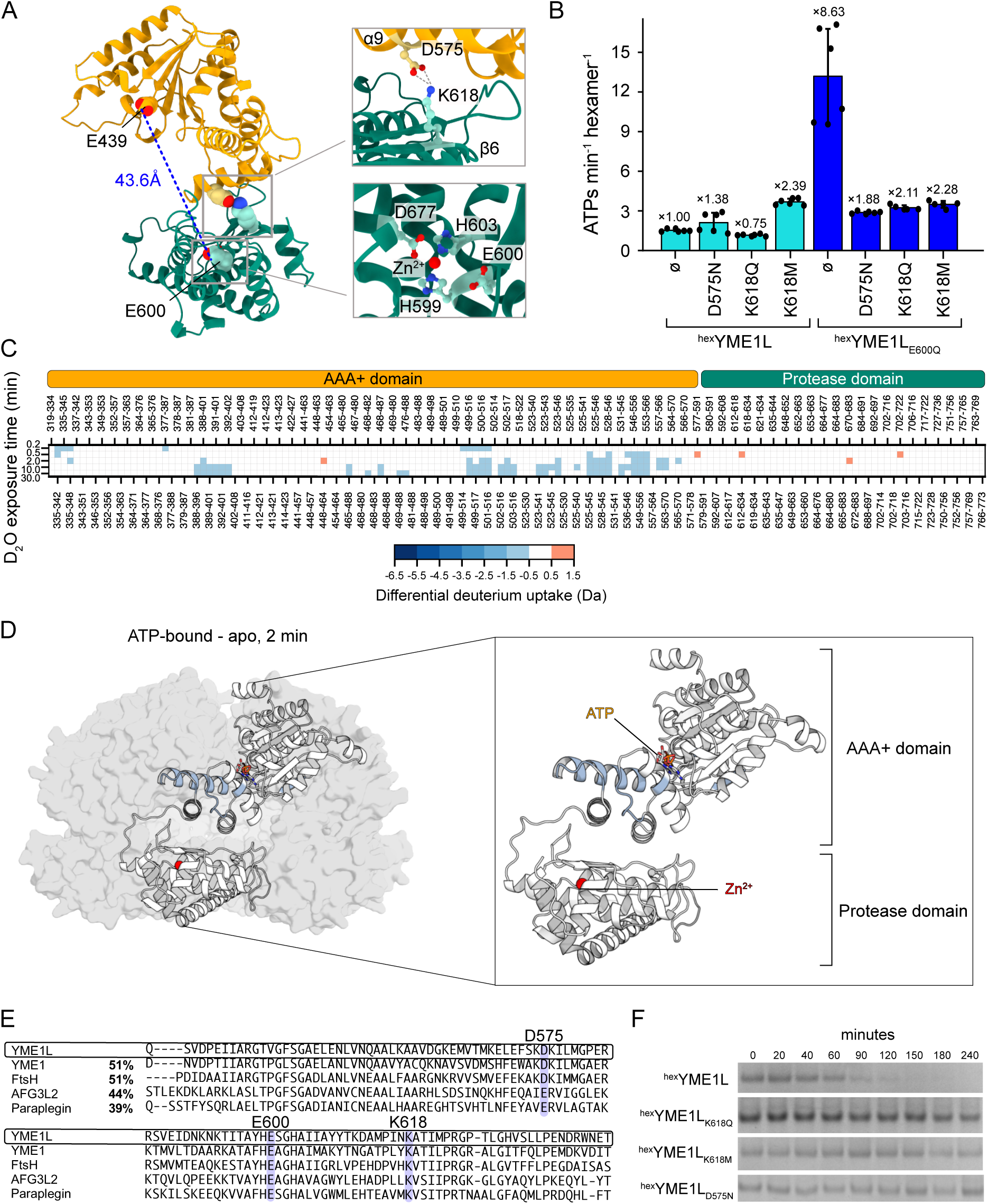
Functional allostery between the AAA+ and the protease domains of ^hex^YME1L. **(A)** Cartoon representation of the catalytic domain of a YME1L subunit predicted by AlphaFold3 (69), highlighting the distance between E439 (Walker-B motif) at the active site of the AAA+ domain and E600 at the catalytic site of the protease domain (left). The domain interface highlighting residues D575 and K618 that are proposed to form a salt bridge and are involved in allosteric communication between the two domains, is enlarged on the upper right. The proteolytic active site is enlarged on the lower right. **(B)** Rates of ATP hydrolysis of ^hex^YME1L mutants, with D575N, K618Q, K618M introduced to either ^hex^YME1L or ^hex^YME1L_E600Q_. Normalized ATPase activities with respect to that of ^hex^YME1L are annotated above the bars. **(C)** Heatmap displays the differential deuterium uptake of ATP-bound ^hex^YME1L_E600Q,E439A_ relative to apo ^hex^YME1L_E600Q,E439A_; **(D)** The differential uptake after 2 minutes of deuterium exposure is mapped onto a YME1L subunit highlighting the rigidification observed in the nucleotide binding site of the AAA+ domain. **(E)** Sequence alignment of YME1L and other homologous AAA+ proteases including *S. cerevisiae* YME1, *E. coli* FtsH, human AFG3L2 and paraplegin. Overall sequence identity of each homolog to YME1L is shown in percentages. Sequence conservation or similarity for D575, E600Q, and K618 across the homologs are highlighted in blue. **(F)** SDS-PAGE showing the lack of degradation of α-casein due to the mutations that disrupt allostery—K618Q, K618M, and D575N, in comparison with that by ^hex^YME1L. Images of the uncropped SDS-PAGE gels are shown in Fig. S12.

**Figure 5.**
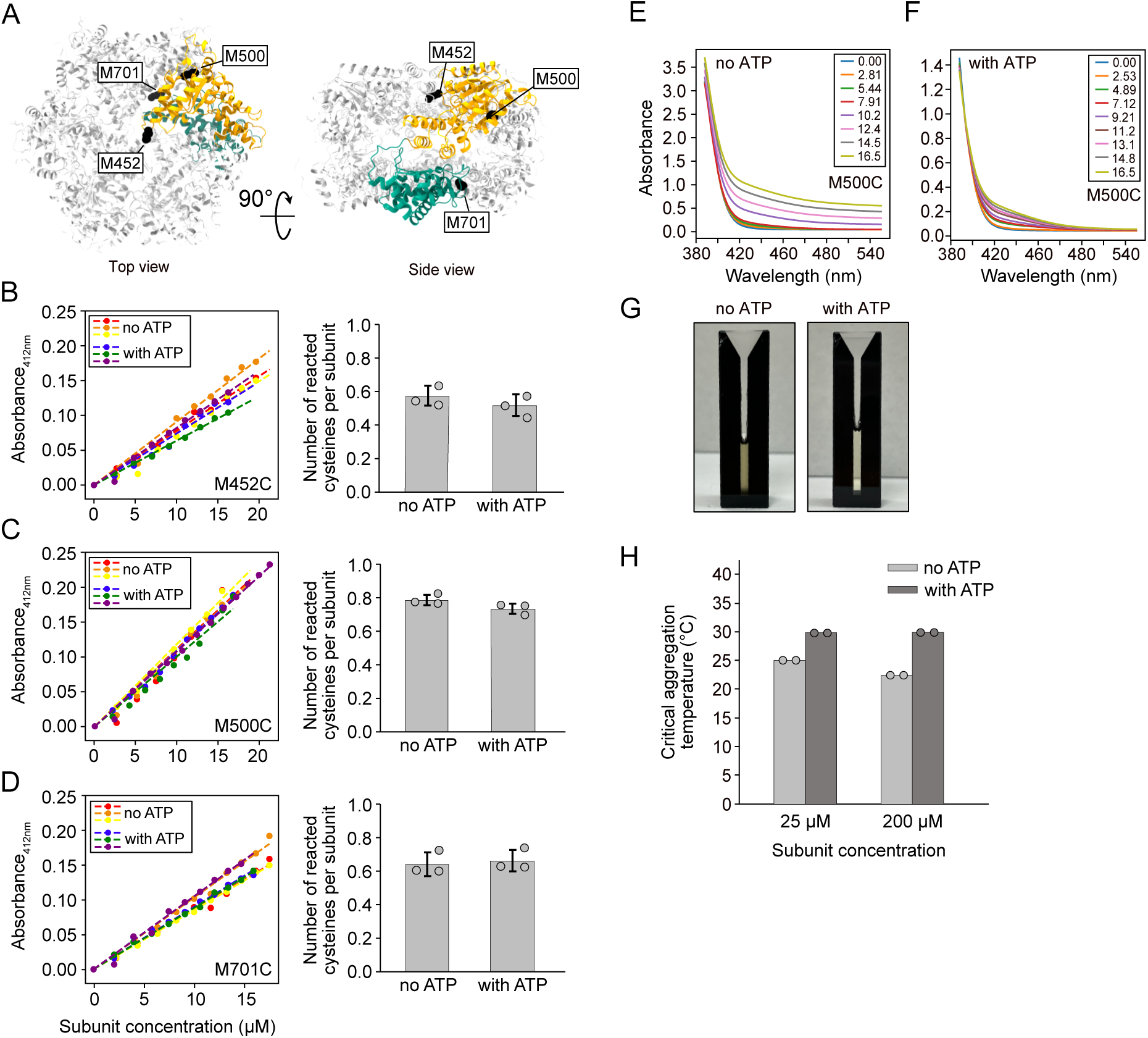
Nucleotide binding increase the stability of YME1L against aggregation. **(A)** Introduction of three cysteine mutations in ^hex^YME1L_E439A_—M452C on the pore-2 loop, M500C and M701C situated at the submit interface of the AAA+ domain and the protease domain, respectively. (B)-(D) Absorbance changes in the DTNB assay (left) with the number of reacted cysteines per subunit (right) for M452C **(B)**, M500C **(C)**, and M701C **(D)**, in the presence and absence of ATP, recorded at 25 °C. The curves on the left are fitted with a linear correlation y = ax. (E-F) Absorbance tracking the titration of M500C into the DTNB solution in the absence **(E)** and presence **(F)** of 5 mM ATP. The concentrations of the ^hex^YME1L subunits are shown in the legend. Aggregation was observed starting at 10.2 μM (subunit concentration) in the absence of ATP, indicated by elevated baselines (E) and turbidity of the solution (**G**, left), while no aggregation occurred in the presence of ATP (F and **G**, right). **(H)** Critical aggregation temperatures of ^hex^YMEL_E439A_ at the subunit concentrations of 25 μM and 200 μM in the presence and absence of 10 mM ATP, measured by DLS.

Upon further examination of the domain interface in the predicted structure of YME1L, we found two residues – D575 (located on α9 in the ATPase domain) and K618 (on β6 in the protease domain), that are within 3 Å of each other, suggesting a possible salt bridge formation (Fig. 4A, upper right). Sequence alignment of YME1L with other homologous AAA+ proteases shows that residues D575 and K618 are conserved (Fig. 4E). To test whether this potential salt bridge is essential for allosteric communication, we introduced point mutations, including D575N, K618Q, and K618M to ^hex^YME1L to disrupt the interaction. Compared to ^hex^YME1L, these point mutations alone resulted in only mild effects on the ATPase activity, with the K618Q mutant showed a 25% decrease, while the D575N and K618M mutants led to 1.38-fold and 2.39-fold increase, respectively (Fig. 4B, teal). However, when these mutations were introduced in the allosterically active ^hex^YME1L_E600Q_ (*i.e.* double mutants of ^hex^YME1L: E600Q & D575N, or E600Q & K618Q, or E600Q & K618M), the ATP hydrolysis rate decreased by approximately 4-fold in comparison to ^hex^YME1L_E600Q_ (Fig. 4B, blue). This indicates that the allosteric activation of the ATPase activity by E600Q is disrupted by mutating the potential salt bridge at the domain interface, suggesting the salt bridge plays an important role in the allosteric communication between the AAA+ and protease domains. We then assessed whether this allosteric communication is essential in the ATP-dependent protease activity of YME1L by carrying out substrate degradation assays on the D575N, K618Q, and K618M mutants of ^hex^YME1L. Despite that all three mutants show comparable or even slightly higher ATPase activity compared to ^hex^YME1L, none of the mutants are able to degrade the model substrate α-casein (Figs. 5E and S12), suggesting the salt bridge between D575 and K618 is crucial for substrate degradation. In summary, these results reveal long-distance functional allostery between the AAA+ and protease domains of YME1L and the importance of a salt bridge between D575 and K618 in bridging the allosteric communication and in facilitating ATP-dependent substrate degradation.

### Nucleotide binding increases the stability of ^hex^YME1L against aggregation

We then used the DTNB assay to investigate whether nucleotide binding affects the solvent accessibility of the subunit interface and the central channel in ^hex^YME1L. DTNB [5,5-dithio-bis-(2-nitrobenzoic acid)] reacts with solvent exposed thiol groups, allowing quantification of reactivity and solvent accessibility of cysteine residues (42). We selected three residues for substitution with cysteines. M500 is located at the subunit interface of the AAA+ domain. M701 is positioned at the subunit interface of the protease domain. M452, in contrast, resides on the pore-2 loop within the central channel (Fig. 5A). The naturally occurring C433 in the ^hex^YME1L construct is not solvent exposed and does not react with DTNB in the assay (Fig. S13). A Walker-B mutation (E439A) was also introduced into the three cysteine variants (M500C, M701C, and M452C) to prevent ATP hydrolysis during the assay. All three YME1L cysteine variants assembles into proper hexamers (Fig. S14). We titrated our ^hex^YME1L variants into a solution with excess DTNB and monitored the absorbance at 412 nm. We quantified the fraction of reacted cysteine side chains in the absence and presence of 5 mM ATP by measuring the slopes of the absorbance change during the titration (Fig. 5B-D). The results revealed that 57%, 79%, and 66% of the cysteines reacted with DTNB in the M452C, M500C and M701C variants, respectively, in the absence of nucleotides, suggesting that all the three positions are partially solvent exposed (Fig. 5B-D, right). This is consistent with the subunit interface and the central channel experiencing a high degree of local dynamics, consistent with our HDX-MS (Fig. 3) and NMR (Fig. 2) data. Surprisingly, we observed minimal differences in solvent accessibility across all mutants in the presence of ATP (Fig. 5B-D). However, increasing concentrations of the TNB-modified ^hex^YME1L led to protein aggregation, indicated by the elevated baselines in the UV-Vis spectra and the apparent turbidity during the assay (Fig. 5E and G, left). Intriguingly, the presence of ATP protects the modified ^hex^YME1L from aggregation at elevated concentrations (Fig. 5F and G, right), potentially by stabilizing the conformation of the AAA+ domain.

In addition to the DTNB analysis, we evaluated the thermal stability of ^hex^YME1L against aggregation using dynamic light scattering (DLS). By monitoring the autocorrelation profiles of ^hex^YME1L as a function of temperature, we leveraged the sensitivity of DLS to detect large aggregates (Fig. S15) (43). Since aggregation is dependent on protein concentrations, we tested ^hex^YME1L at subunit concentrations of 25 and 200 μM. The results show that ^hex^YME1L starts to aggregate when the temperature reaches above 25.0 °C and 22.5 °C (referred to as critical aggregation temperature) in the nucleotide-free state at subunit concentrations of 25 and 200 μM, respectively (Fig. 5H, Fig. S15A and C). Addition of 10 mM ATP, however, increased the thermal stability of ^hex^YME1L, raising the critical aggregation temperatures by 5 °C and 7.5 °C at 25 and 200 μM, respectively (Fig. 5H and Fig. S15B and D). We also observed a similar stabilizing effect of ATP on the monomeric ^ΔN^YME1L construct where ATP binding increases the critical aggregation temperatures of ^ΔN^YME1L at both 25 and 200 μM and protects the protein from thermally induced aggregation (Fig. S16).

## Discussion

Nucleotide binding and hydrolysis are critical components of the functional mechanisms of AAA+ proteins. Oligomerization of the AAA+ domains, often induced or stabilized by nucleotide binding, is essential for their functions, typically by facilitating formation of active sites for ATP hydrolysis at subunit interfaces and the formation of the central channel as the passage of substrates. ATP binding and hydrolysis drives cooperative conformational changes in the subunits, leading to substrate translocation and remodeling (39, 44, 45). Interestingly, for YME1L, nucleotide binding also appears to regulate the degradation of the enzyme itself (22, 24, 46). Previous studies have demonstrated that ATP-independent mitochondrial protease OMA1 rapidly degrades YME1L in response to acute oxidative stress, with ATP depletion being a prerequisite for its degradation. It has been proposed that ATP binding enhances the conformational stability of YME1L, thereby reducing its susceptibility to proteolytic degradation (22, 46). Furthermore, in vitro proteinase K degradation assays have indicated distinct conformational states for YME1L in the presence and absence of nucleotides (24). Although the high-resolution structure of substrate-engaged yeast YME1 has been elucidated (8), the substrate-free conformation of human YME1L (51% homology with the yeast YME1) remains unknown. This raises important questions regarding the mechanism by which nucleotide binding prevents YME1L degradation by the activated OMA1 following mitochondrial membrane depolarization. In this study, we investigated the effects of nucleotide binding on conformational dynamics and structural stability of YME1L using several structural and biophysical tools that include HDX-MS, NMR, dynamic light scattering, and side-chain chemical modification assays. Our findings demonstrate that nucleotide binding stabilizes the catalytic domain of YME1L, providing a potential explanation for how nucleotide binding protects YME1L from OMA1-dependent degradation.

Our HDX-MS data reveal that ATP and ADP binding significantly reduce backbone flexibility in the AAA+ domain of YME1L, particularly within the small helical AAA+ subdomain. Notably, these nucleotide-induced changes in backbone dynamics are observed in both hexameric and monomeric YME1L (Fig. 3 and Fig. S10), suggesting that the rigidification occurs within individual subunits rather than arising from structural rearrangements at subunit interfaces. This observation is consistent with previous structural studies indicating that the large and small AAA+ subdomains undergo substantial movements relative to each other in a nucleotide-dependent manner. These movements coordinate concerted structural rearrangements amongst the subunits during substrate translocation (8). Moreover, our HDX-MS results demonstrate similar deuterium uptake protection patterns for ATP- and ADP-bound states, with the highest protection observed in the ADP-bound state and the lowest in the apo state (Fig. 3C and Fig. S8). Interestingly, prior structural studies indicate that the two AAA+ subdomains are closest in the ADP-bound state and farthest apart in the apo state, coinciding with their relative backbone dynamics at different nucleotide states (8). Additionally, reduced deuterium uptake in the Walker-A motif, which is directly involved in nucleotide binding, has also been observed in other AAA+ motors such as HSP104 (47) and ClpX (48). These findings are consistent with earlier reports showing that binding of nucleotide protects YME1L from hydrogen peroxide-induced conformational changes (24) and prevents its degradation in isolated mitochondria (22). Moreover, ATP binding also mitigates chemical modification-induced and thermally-induced aggregation of YME1L, further emphasizing its structural stabilizing role.

Unlike many AAA+ proteins, truncated YME1L constructs that lack the N-terminal and transmembrane domains exhibit a low intrinsic propensity for oligomerization. Our DLS measurements across a range of protein concentrations, temperatures, and nucleotide conditions show no evidence of oligomerization for the soluble domain. This contrasts with homologous AAA+ proteases, such as FtsH and AFG3L2, that oligomerize either in the absence or presence of nucleotides (26, 49). The low oligomerization propensity of YME1L may result from a loosely packed or highly dynamic subunit interface, which may result in partial solvent accessibility of the engineered cysteines at the subunit interface, as observed in the DTNB assays. The dynamic organization of the hexameric YME1L is also reflected in the poor quality of the NMR spectrum of [^13^CH_3_-ILVM] labeled ^hex^YME1L_AAA_ (Fig. S4A), while improvement of the spectral quality upon the addition of ATPγS suggests that nucleotide binding reduces conformational dynamics and improves conformational homogeneity in the hexamer (Fig. S4B). The dynamic nature of the YME1L catalytic domain likely impedes high-resolution structural determination by cryo-EM in the substrate-free state (8) and could correlate with its rapid degradation under oxidative stress at ATP-depleted conditions (22).

Allostery plays a pivotal role in the functional mechanisms of the AAA+ proteins. In most of classic clade AAA+ proteins (50), ATP binding and hydrolysis are allosterically coupled to the motion of pore loops, which facilitates substrate unfolding and/or translocation (8, 39, 51, 52). Our HDX-MS results reveal reduced deuterium uptake in the pore-2 loop (residues 446-454) upon nucleotide binding, consistent with this conserved translocation mechanism. More interestingly, we observed long-range allosteric communication between the AAA+ and protease domains of YME1L. A mutation in the protease active site leads to a significant increase in the ATPase activity of the enzyme and alters the conformational dynamics of the AAA+ domain. This allosteric communication appears to be mediated by a conserved salt bridge at the domain interface, as mutation of the two residues involved in the salt bridge abolishes allosteric activation of ATPase function and diminishes the proteolytic activity of the enzyme. Long-range allostery, either between different subunits or between the AAA+ and other functional domains, has been reported to play an important role in working mechanisms of AAA+ proteins (26, 51, 53–58). For example, in AAA+ Lon protease, allosteric activation of the proteolytic activity has been reported by substrate binding via induction of a ‘closed’ ring conformation in the AAA+ domain (59). To our knowledge, our results constitute a first observation of a direct coupling between the AAA+ and protease domains in YME1L, contributing to a growing body of evidence that underscores the critical role of interdomain allosteric communication in AAA+ proteins.

In conclusion, our study demonstrates that nucleotide binding not only facilitates YME1L’s enzymatic functions but also plays a pivotal role in modulating its conformational dynamics and structural stability, potentially contributing to the regulation of its degradation under oxidative stress. Furthermore, the observed allosteric communication between nucleotide hydrolysis and proteolytic activity highlights new opportunities to manipulate YME1L function through allosteric mechanisms.

### Experimental procedures

#### Cloning, protein expression and purification

The expression plasmid of ^hex^YME1L from *homo sapiens* in a PET24 vector was a gift from Dr. Justin Miller (Middle Tennessee State University), which encodes a YME1L construct that contains an N-terminal His_6_-tag followed by SUMO (Small Ubiquitin-like Modifier), a hex-tag (GELKAIAQELKAIAKELKAIAWELKAIAQGAG) (6) and the catalytical domain of YME1L (residue 317-773), connected with a 10-aa linker (GSGSYFQSNA). Other YME1L constructs used in the study were generated from the ^hex^YME1L construct using the QuikChange site-directed mutagenesis method with Phusion DNA polymerase (New England Biolabs). The ^hex^YMEL_AAA_ construct contains a hex-tag connected to residues 317-588 of YME1L via the same 10-aa linker. Oligonucleotides used for molecular cloning were synthesized by Integrated DNA Technologies (IDT). Plasmids were prepared by transforming the QuikChange product into the NEB Turbo Competent *E. coli* cells and miniprepped using the NEB Monarch Plasmid Miniprep Kit. All the plasmids were sequenced by Plasmidsaurus to confirm the fidelity of the constructs.

All the YME1L constructs were expressed in *E. coli* strain BL21 (DE3) cells. Cells were grown in the LB broth with 50 μg/mL kanamycin at 37 °C until induction at OD_600_=0.6 with 0.25 mM isopropyl β-d-1-thiogalactopyranoside (IPTG). Protein expression was carried out at 25 °C for *ca.* 16 hours. To express [Ile(δ_1_), Leu(δ_1_,δ_2_), Val(γ_1_,γ_2_), Met(ε), ^13^CH_3_]-labeled YME1L_AAA_ and ^ΔN^YME1L, M9 minimal media in 99% D2O with d_7_-glucose as the sole carbon source was used. Selective methyl labeling was achieved, as previously described (60), by adding 100 mg/L ε-[^13^CH_3_] labeled methionine (CLM-206; Cambridge Isotope Laboratories), 60 mg/L α-ketobutyric acid (CDLM-7318; Cambridge Isotope Laboratories) and 100 mg/L α-ketoisovaleric acid (CDLM-7317; Cambridge Isotope Laboratories) 1 h before IPTG induction at OD_600_ ∼ 0.8. Protein expression was then carried out at 25 °C for *ca.* 16 hours.

Cells were harvested and resuspended in Ni-A buffer [50 mM Tris, pH 7.4, 500 mM NaCl, 50 mM imidazole] followed by lysis via sonication (45% amplitude, 15 min per sample with 2s on/ 2s off pulses). The lysed cells were centrifuged at 14,000 × g for 20 minutes and the supernatant was subjected to 5 mL of HisPur^TM^ Ni-NTA Resin (ThermoScientific). The Ni-NTA Resin column was equilibrated with Ni-A buffer before sample loading, and the bound protein was washed with the Ni-A buffer before elution with Ni-B buffer [25 mM Tris HCl, pH 7.8, 300 mM imidazole]. The eluted protein was then dialyzed overnight at 4 °C in a buffer containing 50 mM Tris (pH7.4), 200 mM NaCl, 1 mM DTT, and ubiquitin-like protease (ULP) to remove the His_6_-SUMO tag. The protein was then concentrated and purified by a Superdex 200 or 75 Increase 10/300 column (Cytiva) in a buffer containing 50 mM HEPES (pH 7.5) and 100 mM NaCl. For the monomeric YME1L constructs, an additional Ni-NTA purification step was used after ULP cleavage to remove the cleaved His_6_-SUMO tag and ULP before the size exclusion chromatography purification step. The purify of all protein preparations was confirmed by SDS-PAGE, and the concentrations were measured using absorbance at 280 nm with extinction coefficients predicted from the primary sequences of the constructs using Expasy ProtParam tool.

The nucleotide-free state of the YME1L constructs were achieved via apyrase digestion. 5 μL of 500 U/ml apyrase (New England BioLabs) was added to the purified YME1L construct in a buffer containing 50 mM HEPES (pH 7.5), 100 mM NaCl, 1 mM EDTA, and 5 mM CaCl_2_, and the mixture was incubated overnight at 21 °C. The apyrase digested sample was purified with Superdex 200 Increase 10/300 column (Cytiva) in a buffer containing 50 mM HEPES (pH 7.5), 100 mM NaCl, and 1 mM EDTA. The fractions of pure protein were collected, flash frozen in liquid nitrogen, and stored at −80 °C.

#### Size-exclusion chromatography

To characterize the YME1L constructs in the presence of ATP using size-exclusion chromatography (SEC), 30 μM of purified ^ΔN^YME1L_E439Q_ and ^hex^YME1L_E439A_ were first incubated in a buffer containing 50 mM HEPES (pH 7.5), 100 mM NaCl, 2 mM ATP, 5 mM MgCl_2_, and 1 mM EDTA for 2 hours prior to SEC runs. A total volume of 0.3 mL protein samples was then injected onto a Superdex 200 Increase 10/300 column (Cytiva) equilibrated with a buffer containing 50 mM HEPES (pH 7.5), 100 mM NaCl, 1 mM EDTA, 2 mM ATP, and 5 mM MgCl_2_.

#### Dynamic light scattering

Purified ^hex^YME1L_E439A_ were stored in the apo form. For the ATP-bound samples, 10 mM ATP was added to the protein solution prior to the DLS experiments. DLS measurements were performed on a DynaPro DLS Plate Reader III (Wyatt Technology) in a buffering containing 50 mM HEPES (pH 7.5), 100 mM NaCl, 1 mM EDTA, 15 mM MgCl_2_. Samples were centrifuged at 20,000 × g at 4 °C for 30 min to eliminate precipitates or solid impurities prior to loading on a 384-well plate. A total of 20 μL of protein sample was loaded to each well, and the plate was centrifuged at 3,000 × g at 4 °C for 5 minutes to remove air bubbles present in the wells. 10 μL of mineral oil was then added on top of each sample to prevent evaporation, and the plate was centrifuged again at 3,000 × g centrifugation at 4 °C for 3 minutes. For each well, 20 measurements with 4.8-s acquisitions were performed and an average was calculated from the autocorrelation functions. Temperature scans were performed from 7.5 °C to 50 °C with a step size of 2.5 °C. The autocorrelation functions were fitted using the following equation

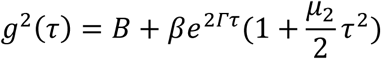

where *B* represents a term accounting for the baseline, *β* is the correlation function amplitude at τ=0, Γ = Dq^2^, and μ_2_ is the second moment of the distribution of Γ values about their mean (61, 62). For the calculation of Γ, *D* is the translational diffusion coefficient and *q* is the scattering wave vector calculated by 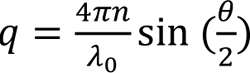, where *n* is the solvent refractive index (*n*=1.3347), λ_0_ is the wavelength of the light used by the instrument, and *θ* is the scattering angle. *D*, *β* and *B* and μ_2_ were output as fitted parameters, respectively.

#### NADH-coupled ATPase assay

NADH-coupled ATPase assays (25) were carried out at 25 °C in a reaction buffer containing 25 mM HEPES (pH 7.5), 5 mM MgCl_2_, 100 mM NaCl, 257 μM NADH, 50 U/ml LDH, and an ATP regeneration system (2 mM ATP, 1 mM PEP, and 20 U/mL PK). A total reaction volume of 100 μL was used with 1.8 μM (hexamer concentration) of YME1L construct in each reaction. To measure the ATP hydrolysis rate in the presence of Zn^2+^, a final concentration of 25 μM ZnCl_2_ was included in the reaction. Reactions were carried out in triplicates. The decrease in absorbance at 340 nm was monitored in a 96-well plate using an Epoch 2 Microplate Spectrophotomer (BioTek Instruments). The ATPase hydrolysis rate was calculated by using Beer’s law and the NADH extinction coefficient at 340 nm (ε_340_ = 6,220 M^-1^cm^-1^).

#### Protein degradation assays

Substrate degradation assays were conducted at 29 °C using 2.6 μM (hexamer concentration) of YME1L in a buffer containing 25 mM HEPES (pH 7.5), 100 mM NaCl, 10 mM MgCl_2_, 1 mM DTT, 25 μM ZnCl_2_, and an ATP regeneration system (5 mM ATP, 2 mM PEP, and 2.5 U mL^-1^ PK). 10 μM of α-casein (Sigma Aldrich, Catalogue #C6780) was used in the reaction. Aliquots were sampled from the reaction mix at the various time points and mixed with a 2× SDS-loading buffer (FroggaBio Inc.) including 2 mM DTT to quench the reaction. Degradation was evaluated by SDS-PAGE to separate the band followed by Coomassie Brilliant Blue G-250 staining. The SDS-PAGE gels were imaged using the BioRad ChemiDocTsM XRS+ gel documentation system, and the band intensities of α-casein as a function of reaction time were quantified using Image LabTM software (Bio-Rad).

#### NMR experiments

All protein samples were buffer exchanged using centrifugal filters (Amicon Ultra, 10 kDa MWCO, Millipore Sigma) into D_2_O NMR buffer [25mM HEPES, pH 7.0 (pD 7.4), 25mM NaCl, 1mM EDTA]. The protein samples were centrifuged at 6,000 ×g for 1 minute to remove any aggregates prior to NMR experiments. NMR experiments were conducted on Bruker AVANCE III 14.1 T spectrometer, equipped with cryogenically cooled, pulse-field gradient, triple-resonance probes. The NMR spectra collected were processed with NMRPipe (66) and visualized using NMRFAM-SPARKY (67). The ^1^H-^13^C HMQC experiments were acquired with an interscan delay of 1.5 s with the carriers positioned at 19 ppm and 0.8 ppm and acquisition times of 21.3 ms and 64 ms at the t_1_ and t_2_ dimensions, respectively.

In order to measure the binding affinity of ADP to the monomeric YME1L_AAA_, a titration was performed in which ADP was added to a solution of ILVM-methyl labeled AAA+ domain (maintained at 100 μM) in a series of 13 steps from 0 to 392 μM. Assuming a one-site binding model, the binding curves were individually fit to the following equation using nonlinear least square fit:

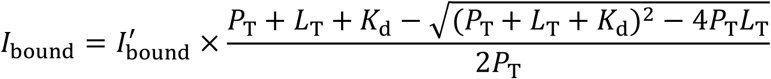

where *I*_bound_ is the intensity of the bound peak at each titration point, *I*^′^ is the intensity of the bound peak at the fully bound state, *P*_T_ and *L*_T_ are the total concentrations of YME1L_AAA_ and ADP, respectively. Fitting was performed using an in-house Python script employing the ‘lmfit’ package with *K*_d_ and *I*^′^ being the fitted parameters. The final *K*_d_ is the average of the individually fitted *K*_d_ values with uncertainty obtained from the standard deviation.

#### HDX-MS

To prepare the samples for the HDX-MS experiments, protein stocks were diluted into an equilibration buffer in H_2_O containing 50 mM HEPES (pH 7.5), 100 mM NaCl, 1 mM EDTA, and 10 mM MgCl_2_. The buffers were supplemented with 10 mM ATP or ADP to create the ATP-bound and ADP-bound states, respectively. For the (ATP, Zn^2+^)-bound ^hex^YME1L, the protein was incubated in 50 mM HEPES (pH 7.5), 100 mM NaCl, 10 mM MgCl_2_, 10 mM ATP and 1 mM of ZnCl_2_. HDX reaction was initiated by a ten-fold dilution into a D_2_O-based buffer with the same composition as the equilibration buffer and additives depending on the condition of interest at pD 7.5 (63). HDX reactions were quenched at fixed time points of 0.167, 0.5, 2, 10, 30 minutes after the exchange process via acidification to pH of 2.5 in a 1:1 ratio of HDX reaction and quench buffer (250 mM Na_3_PO_4_, 3 M Gdn-HCl, 3 mM MAPCHO-12, pH 1.37), followed by flash freezing in liquid N_2_. The final sample contained 0.033 μM (hexamer concentration) YME1L with a D_2_O concentration of ∼90%. Samples were stored at −80°C until LC-MS analysis. All time points and conditions were run in triplicate to assess the reproducibility of the measurements.

For LC-MS, the samples were rapidly thawed and injected into the 50 μL loop of an M-class ACQUITY ultra-performance liquid chromatography (UPLC) equipped with HDX technology (Waters). The protein was digested into smaller peptide fragments via an immobilized Nepenthesin II column (AP-PC-004m 1 mm x 20 mm, 16.2 μL vol., AffiPro). The generated peptic peptides were trapped on a C18 Vanguard BEH column (ACQUITY UPLC BEH, 1.7 μM, 2.1 mm x 5 mm, Waters) for 3 minutes at 100 μL min^-1^ prior to MS analysis. While maintaining 0 °C, trapped peptides were subsequently separated on a UPLC HSS T3 column (ACQUITY, 1.8 μM, 1 mm x 50 mm, Waters) at 100 μL min^-1^ rate with an 8-minute water/acetonitrile gradient acidified with 0.1% formic acid. To minimize the carryover of samples, the trapping and analytical columns were cleaned for 12 extra minutes. Additionally, pepsin wash (1.5 M Gdn-HCl, 5% acetonitrile,

0.8% formic acid, 1.5 mM MAPCHO-12) was injected 5 times during each run. The LC eluent was directed to a Synapt G2Si mass spectrometer (Waters) equipped with a dual electrospray ionization source (ESI) operated in positive-ion mode. The mass spectrometer fitted with a standard ESI was operated in positive ion mode with a capillary voltage of +3 kV. The instrument was calibrated online via infusion of LeuEnk solution (1+, 556.2771 *m/z*) from the LockSpray exact mass ionization source (Waters) every 20 seconds at a flow rate of 10 μL min^-1^. The cone and desolvation gas flows were set to 90 and 600 L hr ^-1^, respectively. The source and desolvation temperatures were 80 and 175 °C, respectively. The instrument was operated in resolution mode with an acquisition *m/z* range of 50 to 2000 Th. Subsequently, HDX data analysis to measure deuterium uptake was performed using DynamX 3.0 software (Waters).

Peptide mapping was performed in triplicate on undeuterated samples using data-independent acquisition (DIA), also known as MS^E^ mode. The MS^E^ datasets were processed by ProteinLynx Global Server (PLGS) software program (Waters) searching against a database of the YME1L construct sequence. Additionally, the PLGS processing parameters were set to lock mass window of 0.4 Da, low energy and elevated energy threshold of 100 and 50 counts, respectively. PLGS generated ion accounting files in CSV format consisting of a list of all identified peptides for all three runs of the undeuterated control samples. Information including individual peptide sequences, the number of matched product ions, the charge number, *m/z* value of all precursor ions were provided from the ion accounting files. Subsequently, YME1L peptides that were identified in all triplicates after filtering using the parameters suggested by Sørensen and colleagues were included to be processed further (64).

The identified peptides from PLGS were subsequently analyzed using DynamX 3.0 software. The mass spectra of all the peptides under each exchange time point were manually checked to maximize the analysis quality and overall sequence coverage. The deuterium uptake profiles, plotting the deuterium uptake of individual exposure points in all states for individual peptides were generated once the mass spectra were fully analyzed. The centroid mass of both deuterated and undeuterated peptide isotope distribution under each time point was obtained according to this equation:

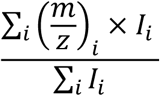

where the spectral intensity of *i*^th^ channel in the spectrum is represented by I_i_ (65). The deuterium uptake differences between conditions were calculated by comparing the centroids of isotope distribution between the deuterated and undeuterated samples. HDX-MS results were also visualized as heat maps to display the deuterium uptake differences across states and protein constructs in a comparative fashion. The experimental details of HDX-MS were summarized in Table S1.

#### DTNB assay

Prior to the DTNB assay, the cysteine mutants were incubated with 10 mM DTT to reduce the cysteine followed by size exclusion chromatography to remove DTT from the protein stock solution. 10 mM DTNB was prepared in a buffer containing 100 mM sodium phosphate (pH 7.8) and then diluted to 1 mM in a buffer containing 50 mM HEPES (pH 7.5), 100 mM NaCl, and 1 mM EDTA. A titration was performed by adding 10 μL protein (80∼200 μM monomeric concentration) consecutively to the 1 mM DTNB solution and monitor the reaction using UV-Vis absorbance. For the ATP-bound state, a final concentration of 5 mM ATP and 10 mM MgCl_2_ was included in both the DTNB and the protein solution prior to the titration. The absorbance at 412 nm was plotted as a function of protein concentration and fitted with a linear function (*y* = *kx*), in which the slope was used to extract the fraction of reacted thiol groups using an extinction coefficient of 14150 cm^-1^ M^-1^ for TNB (68).

## Data availability

All processed data are available in the article and supporting information. Mass spectrometry data are available from the MassIVE database under entry MSV000096916.

## Supporting information

This article contains supporting information.

## Acknowledgments

M.K.B. acknowledges support for an NSERC Canada Graduate Scholarship. C.Y.C. and M.M.G. acknowledge support for Ontario Graduate Scholarship. Financial support was provided by a Natural Sciences and Engineering Research Council of Canada Discovery Grant (RGPIN-2021-02843) to S.V. and (RGPIN-2020-00527) to R.H.. MS data were recorded at the Mass Spectrometry Facility of the Advanced Analysis Centre, University of Guelph. We thank Dr. Dyanne Brewer (University of Guelph) for assistance with MS measurements. NMR data were recorded at the University of Guelph NMR Center. We thank Dr. Sameer Al-Abdul-Wahid (University of Guelph) for the support with NMR measurements. The DLS data were recorded in the Structural & Biophysical Core Facility at the Hospital for Sick Children, Toronto. We thank Dr. Robert W. Harkness (University of Guelph) for the assistance with the data acquisition. We thank Dr. Algirdas Velyvis for critical review of the manuscript.

## Author contributions

M. K. B., A. S. K., C. Y. C., M. M. G., S. W., S. V., and R. H. formal analysis; M. K. B., A. S. K., C. Y. C., M. M. G., S. W., S. V., and R. H. investigation; S. V. and R. H. supervision; M. K. B., A. S. K., C. Y. C., M. M. G., S. V., and R. H. visualization; M. K. B., A. S. K., S. V., and R. H. writing – original draft; M. K. B., M. M. G., S. V., and R. H. writing–review & editing; M. K. B., A. S. K., C. Y. C., M. M. G., S. W., S. V., and R. H. methodology; S. V. and R. H. resources; S. V. and R. H. funding acquisition, S. V. and R. H. project administration.

## Abbreviations

AAA+: ATPases Associated with diverse cellular Activities

ADP: adenosine diphosphate

ATP: adenosine triphosphate

ATPγS: adenosine 5′-O-(3-thio)triphosphate

DLS: dynamic light scattering

DTNB: 5,5-dithio-bis-(2-nitrobenzoic acid)

DTT: dithiothreitol

ESI: electrospray ionization

Gdn-HCl: guanidine hydrochloride

HDX: hydrogen deuterium exchange

IPTG: isopropyl-β-D-1-thiogalactopyranoside

IMM: inner mitochondrial membrane

IMS: intermembrane space

*K*_d_: equilibrium dissociation constant

LC: liquid chromatography

LDH: lactate dehydrogenase

L-OPA1: long form of OPA1 m/z – mass to charge ratio

MS: mass spectrometry

MWCO: molecular weight cut-off

NMR: nuclear magnetic resonance

NADH: reduced nicotinamide adenine dinucleotides

NTD: N-terminal domain

OD_600_: optical density

PAGE: polyacrylamide gel electrophoresis

PEP: phosphoenolpyruvate

PK: pyruvate kinase

SDS: sodium dodecyl sulphate

SEC: size exclusion chromatography

S-OPA1: shot form of OPA1

SUMO: small ubiquitin-like modifier

TMD: transmembrane domain

TNB: 2-nitro-5-thiobenzoate

TROSY: transverse relaxation optimized spectroscopy

ULP: ubiquitin-like protease

UPLC: ultra-performance liquid chromatography

UV: ultraviolet

Vis: visible

## References

1. Pagliarini, D. J., Calvo, S. E., Chang, B., Sheth, S. A., Vafai, S. B., Ong, S.-E., Walford, G. A., Sugiana, C., Boneh, A., Chen, W. K., Hill, D. E., Vidal, M., Evans, J. G., Thorburn, D. R., Carr, S. A., and Mootha, V. K. (2008) A Mitochondrial Protein Compendium Elucidates Complex I Disease Biology. Cell. 134, 112–123

2. Lopez, M. F., Kristal, B. S., Chernokalskaya, E., Lazarev, A., Shestopalov, A. I., Bogdanova, A., and Robinson, M. (2000) High-throughput profiling of the mitochondrial proteome using affinity fractionation and automation. Electrophoresis. 21, 3427–3440

3. Song, J., Herrmann, J. M., and Becker, T. (2021) Quality control of the mitochondrial proteome. Nat Rev Mol Cell Biol. 22, 54–70

4. Steele, T. E., and Glynn, S. E. (2019) Mitochondrial AAA proteases: A stairway to degradation. Mitochondrion. 49, 121–127

5. Feng, Y., Nouri, K., and Schimmer, A. D. (2021) Mitochondrial ATP-Dependent Proteases—Biological Function and Potential Anti-Cancer Targets. Cancers (Basel). 13, 2020

6. Shi, H., Rampello, A. J., and Glynn, S. E. (2016) Engineered AAA+ proteases reveal principles of proteolysis at the mitochondrial inner membrane. Nat Commun. 7, 13301

7. Coppola, M., Pizzigoni, A., Banfi, S., Bassi, M. T., Casari, G., and Incerti, B. (2000) Identification and Characterization of YME1L1, a Novel Paraplegin-Related Gene. Genomics. 66, 48–54

8. Puchades, C., Rampello, A. J., Shin, M., Giuliano, C. J., Wiseman, R. L., Glynn, S. E., and Lander, G. C. (2017) Structure of the mitochondrial inner membrane AAA+ protease YME1 gives insight into substrate processing. Science (1979). 358, eaao0464

9. Rainbolt, T. K., Saunders, J. M., and Wiseman, R. L. (2015) YME1L degradation reduces mitochondrial proteolytic capacity during oxidative stress. EMBO Rep. 16, 97–106

10. Qi, Y., Liu, H., Daniels, M. P., Zhang, G., and Xu, H. (2016) Loss of Drosophila i-AAA protease, dYME1L, causes abnormal mitochondria and apoptotic degeneration. Cell Death & Differentiation. 23, 291–302

11. Abass, S. A., Abdel-Hamid, N. M., Elshazly, A. M., Abdo, W., and Zakaria, S. (2023) OMA1 and YME1L as a Diagnostic Panel in Hepatocellular Carcinoma. Yale J Biol Med. 96, 443– 454

12. Sun, X., Shi, C., Dai, J., Zhang, M.-Q., Pei, D.-S., and Yang, L. (2024) Targeting the mitochondrial protein YME1L to inhibit osteosarcoma cell growth in vitro and in vivo. Cell Death Dis. 15, 346

13. Xia, Y., He, C., Hu, Z., Wu, Z., Hui, Y., Liu, Y.-Y., Mu, C., and Zha, J. (2023) The mitochondrial protein YME1 Like 1 is important for non-small cell lung cancer cell growth. Int J Biol Sci. 19, 1778–1790

14. Cheng, F., Huang, H., Yin, S., Liu, J.-S., and Sun, P. (2024) Expression and functional implications of YME1L in nasopharyngeal carcinoma. Cell Death Dis. 15, 423

15. Stiburek, L., Cesnekova, J., Kostkova, O., Fornuskova, D., Vinsova, K., Wenchich, L., Houstek, J., and Zeman, J. (2012) YME1L controls the accumulation of respiratory chain subunits and is required for apoptotic resistance, cristae morphogenesis, and cell proliferation. Mol Biol Cell. 23, 1010–1023

16. Macvicar, T., Ohba, Y., Nolte, H., Mayer, F. C., Tatsuta, T., Sprenger, H.-G., Lindner, B., Zhao, Y., Li, J., Bruns, C., Krüger, M., Habich, M., Riemer, J., Schwarzer, R., Pasparakis, M., Henschke, S., Brüning, J. C., Zamboni, N., and Langer, T. (2019) Lipid signalling drives proteolytic rewiring of mitochondria by YME1L. Nature. 575, 361–365

17. Anand, R., Wai, T., Baker, M. J., Kladt, N., Schauss, A. C., Rugarli, E., and Langer, T. (2014) The i-AAA protease YME1L and OMA1 cleave OPA1 to balance mitochondrial fusion and fission. Journal of Cell Biology. 204, 919–929

18. Del Dotto, V., Mishra, P., Vidoni, S., Fogazza, M., Maresca, A., Caporali, L., Mccaffery, J. M., Cappelletti, M., Baruffini, E., Lenaers, G., Chan, D., Rugolo, M., Carelli, V., and Zanna, C. (2017) OPA1 Isoforms in the Hierarchical Organization of Mitochondrial Functions. Cell Rep. 19, 2557–2571

19. Ban, T., Ishihara, T., Kohno, H., Saita, S., Ichimura, A., Maenaka, K., Oka, T., Mihara, K., and Ishihara, N. (2017) Molecular basis of selective mitochondrial fusion by heterotypic action between OPA1 and cardiolipin. Nat Cell Biol. 19, 856–863

20. Zhang, D., Zhang, Y., Ma, J., Zhu, C., Niu, T., Chen, W., Pang, X., Zhai, Y., and Sun, F. (2020) Cryo-EM structures of S-OPA1 reveal its interactions with membrane and changes upon nucleotide binding. Elife. 9, e50294

21. 21. Van Der Bliek, A. M., Shen, Q., and Kawajiri, S. (2013) Mechanisms of Mitochondrial Fission and Fusion. Cold Spring Harb Perspect Biol. 5, a011072

22. Zhang, K., Li, H., and Song, Z. (2014) Membrane depolarization activates the mitochondrial protease OMA1 by stimulating self-cleavage. EMBO Rep. 15, 576–585

23. Wai, T., García-Prieto, J., Baker, M. J., Merkwirth, C., Benit, P., Rustin, P., Rupérez, F. J., Barbas, C., Ibañez, B., and Langer, T. (2015) Imbalanced OPA1 processing and mitochondrial fragmentation cause heart failure in mice. Science (1979). 350, aad0116

24. Brambley, C. A., Marsee, J. D., Halper, N., and Miller, J. M. (2019) Characterization of Mitochondrial YME1L Protease Oxidative Stress-Induced Conformational State. J Mol Biol. 431, 1250–1266

25. Norby, J. G. (1988) Coupled Assay of Na+,K+-ATPase Activity. Methods Enzymol. 156, 116–119

26. Puchades, C., Ding, B., Song, A., Wiseman, R. L., Lander, G. C., and Glynn, S. E. (2019) Unique Structural Features of the Mitochondrial AAA+ Protease AFG3L2 Reveal the Molecular Basis for Activity in Health and Disease. Mol Cell. 75, 1073–1085.e6

27. Li Binz, T., Niemann, H., and Singh, B. R. (2000) Probing the Mechanistic Role of Glutamate Residue in the Zinc-Binding Motif of Type A Botulinum Neurotoxin Light Chain. Biochemistry. 39, 2399–2405

28. Rosenzweig, R., and Kay, L. E. (2016) Solution NMR Spectroscopy Provides an Avenue for the Study of Functionally Dynamic Molecular Machines: The Example of Protein Disaggregation. J Am Chem Soc. 138, 1466–1477

29. Tugarinov, V., Hwang, P. M., Ollerenshaw, J. E., and Kay, L. E. (2003) Cross-correlated relaxation enhanced 1H-13C NMR spectroscopy of methyl groups in very high molecular weight proteins and protein complexes. J Am Chem Soc. 125, 10420–10428

30. Jiang, Y., and Kalodimos, C. G. (2017) NMR Studies of Large Proteins. J Mol Biol. 429, 2667–2676

31. Bain, A. D. (2003) Chemical exchange in NMR. Prog Nucl Magn Reson Spectrosc. 43, 63– 103

32. Palmer, A. G., Kroenke, C. D., and Patrick Loria, J. (2001) [10] - Nuclear Magnetic Resonance Methods for Quantifying Microsecond-to-Millisecond Motions in Biological Macromolecules. in Methods in Enzymology (James, T. L., Dötsch, V., and Schmitz, U. eds), pp. 204–238, Academic Press, 339, 204–238

33. Miller, J. M., Brambley, C. A., and Marsee, J. D. (2020) Examination of the Role of Mg2+in the Mechanism of Nucleotide Binding to the Monomeric YME1L AAA+ Domain. Biochemistry. 59, 4303–4320

34. Skinner, J. J., Lim, W. K., Bédard, S., Black, B. E., and Englander, S. W. (2012) Protein hydrogen exchange: Testing current models. Protein Science. 21, 987–995

35. Ye, X., Lin, J., Mayne, L., Shorter, J., and Englander, S. W. (2020) Structural and kinetic basis for the regulation and potentiation of Hsp104 function. PNAS. 117, 9384–9392

36. Konermann, L., Rodriguez, A. D., and Sowole, M. A. (2014) Type 1 and Type 2 scenarios in hydrogen exchange mass spectrometry studies on protein–ligand complexes. Analyst. 139, 6078–6087

37. Hanson, P. I., and Whiteheart, S. W. (2005) AAA+ proteins: have engine, will work. Nat Rev Mol Cell Biol. 6, 519–529

38. Sauer, R. T., and Baker, T. A. (2011) AAA+ Proteases: ATP-Fueled Machines of Protein Destruction. Annurev Biochem. 80, 587–612

39. Puchades, C., Sandate, C. R., and Lander, G. C. (2020) The molecular principles governing the activity and functional diversity of AAA+ proteins. Nat Rev Mol Cell Biol. 21, 43–58

40. Banerjee, S., Bartesaghi, A., Merk, A., Rao, P., Bulfer, S. L., Yan, Y., Green, N., Mroczkowski, B., Neitz, R. J., Wipf, P., Falconieri, V., Deshaies, R. J., Milne, J. L. S., Huryn, D., Arkin, M., and Subramaniam, S. (2016) 2.3 Å resolution cryo-EM structure of human p97 and mechanism of allosteric inhibition. Science. 351, 871–875

41. Gerdes, F., Tatsuta, T., and Langer, T. (2012) Mitochondrial AAA proteases - Towards a molecular understanding of membrane-bound proteolytic machines. Biochim Biophys Acta Mol Cell Res. 1823, 49–55

42. Ellman, G. L. (1959) Tissue Sulfhydryl Groups. Arch Biochem Biophys. 82, 70–77

43. Some, D. (2013) Light-scattering-based analysis of biomolecular interactions. Biophys Rev. 5, 147–158

44. Gates, S. N., and Martin, A. (2020) Stairway to translocation: AAA+ motor structures reveal the mechanisms of ATP-dependent substrate translocation. Protein Science. 29, 407–419

45. Khan, Y. A., White, K. I., and Brunger, A. T. (2022) The AAA+ superfamily: a review of the structural and mechanistic principles of these molecular machines. Crit Rev Biochem Mol Biol. 57, 156–187

46. T, Lebeau, J., Puchades, C., and R (2016) Reciprocal Degradation of YME1L and OMA1 Adapts Mitochondrial Proteolytic Activity during Stress. Cell Rep. 14, 2041–2049

47. Ye, X., Lin, J., Mayne, L., Shorter, J., and Englander, S. W. (2019) Hydrogen exchange reveals Hsp104 architecture, structural dynamics, and energetics in physiological solution. PNAS. 116, 7333–7342

48. Feng, Y., Goncalves, M. M., Jitkova, Y., Keszei, A., Sarathy, C., Trudel, V., St-Germain, J., Tcheng, M., Yan, Y., Hurren, R., Schultz, M., Raught, B., Yudin, A., Mazhab-Jafari, M., Vahidi, S., and Schimmer, A. D. (2024) Abstract 7053: Serine phosphorylation marks proteins for degradation by the mitochondrial matrix protease, ClpXP. Cancer Res. 84, 7053

49. Bieniossek, C., Niederhauser, B., and Baumann, U. M. (2009) The crystal structure of apo-FtsH reveals domain movements necessary for substrate unfolding and translocation. PNAS. 106, 21579–21584

50. Seraphim, T. V, and Houry, W. A. (2020) AAA+ proteins. Curr Biol. 30, R251–R257

51. Pan, M., Yu, Y., Ai, H., Zheng, Q., Xie, Y., Liu, L., and Zhao, M. (2021) Mechanistic insight into substrate processing and allosteric inhibition of human p97. Nat Struct Mol Biol. 28, 614–625

52. Iljina, M., Mazal, H., Dayananda, A., Zhang, Z., Stan, G., Riven, I., and Haran, G. (2024) Single-molecule FRET probes allosteric effects on protein-translocating pore loops of a AAA+ machine. Biophys J. 123, 374–388

53. Augustin, S., Gerdes, F., Lee, S., Tsai, F. T. F., Langer, T., and Tatsuta, T. (2009) An Intersubunit Signaling Network Coordinates ATP Hydrolysis by m-AAA Proteases. Mol Cell. 35, 574–585

54. Caffrey, B., Zhu, X., Berezuk, A., Tuttle, K., Chittori, S., and Subramaniam, S. (2021) AAA+ ATPase p97/VCP mutants and inhibitor binding disrupt inter-domain coupling and subsequent allosteric activation. J Biol Chem. 297, 101187

55. Huang, R., Ripstein, Z. A., Rubinstein, J. L., and Kay, L. E. (2019) Cooperative subunit dynamics modulate p97 function. PNAS. 116, 158–167

56. Mickolajczyk, K. J., Olinares, P. D. B., Niu, Y., Chen, N., Warrington, S. E., Sasaki, Y., Walz, T., Chait, B. T., and Kapoor, T. M. (2020) Long-range intramolecular allostery and regulation in the dynein-like AAA protein Mdn1. PNAS. 117, 18459–18469

57. Mazal, H., Iljina, M., Barak, Y., Elad, N., Rosenzweig, R., Goloubinoff, P., Riven, I., and Haran, G. (2019) Tunable microsecond dynamics of an allosteric switch regulate the activity of a AAA+ disaggregation machine. Nat Commun. 10, 1438

58. Shein, M., Hitzenberger, M., Cheng, T. C., Rout, S. R., Leitl, K. D., Sato, Y., Zacharias, M., Sakata, E., and Schütz, A. K. (2024) Characterizing ATP processing by the AAA+ protein p97 at the atomic level. Nat Chem. 16, 363–372

59. Shin, M., Puchades, C., Asmita, A., Puri, N., Adjei, E., Wiseman, R. L., Karzai, A. W., and Lander, G. C. (2020) Structural basis for distinct operational modes and protease activation in AAA+ protease Lon. Sci Adv. 6, eaba8404

60. Conicella, A. E., Huang, R., Ripstein, Z. A., Nguyen, A., Wang, E., Löhr, T., Schuck, P., Vendruscolo, M., Rubinstein, J. L., and Kay, L. E. (2020) An intrinsically disordered motif regulates the interaction between the p47 adaptor and the p97 AAA+ ATPase. Proc Natl Acad Sci U S A. 117, 26226–26236

61. Koppel, D. E. (1972) Analysis of macromolecular polydispersity in intensity correlation spectroscopy: The method of cumulants. J Chem Phys. 57, 4814–4820

62. Frisken, B. J. (2001) Revisiting the method of cumulants for the analysis of dynamic light-scattering data. Appl. Opt. 10, 4087–4091

63. Krȩżel, A., and Bal, W. (2004) A formula for correlating pKa values determined in D2O and H2O. J Inorg Biochem. 98, 161–166

64. Sørensen, L., and Salbo, R. (2018) Optimized Workflow for Selecting Peptides for HDX-MS Data Analyses. J Am Soc Mass Spectrom. 29, 2278–2281

65. Weis, D. D., Engen, J. R., and Kass, I. J. (2006) Semi-Automated Data Processing of Hydrogen Exchange Mass Spectra Using HX-Express. J Am Soc Mass Spectrom. 17, 1700– 1703

66. Delaglio, F., Grzesiek, S., Vuister, G. W., Zhu, G., Pfeifer, J., and Bax, A. (1995) NMRPipe: A multidimensional spectral processing system based on UNIX pipes. J Biomol NMR. 6, 277–293

67. Lee, W., Tonelli, M., and Markley, J. L. (2015) NMRFAM-SPARKY: enhanced software for biomolecular NMR spectroscopy. Bioinformatics. 31, 1325–1327

68. Riddles, P. W., Blakeley, R. L., and Zerner, B. Ellman’s Reagent: 5,5’-Dithiobis(2-nitrobenzoic Acid)-a Reexamination. Anal Biochem. 94, 75–81

69. Abramson, J., Adler, J., Dunger, J., Evans, R., Green, T., Pritzel, A., Ronneberger, O., et al. (2024) Accurate structure prediction of biomolecular interactions with AlphaFold 3. Nature. 630, 493–500

