## Supplementary Figures for "Characterization of conformational dynamics and allostery of the catalytic domain of human mitochondrial YME1L protease"

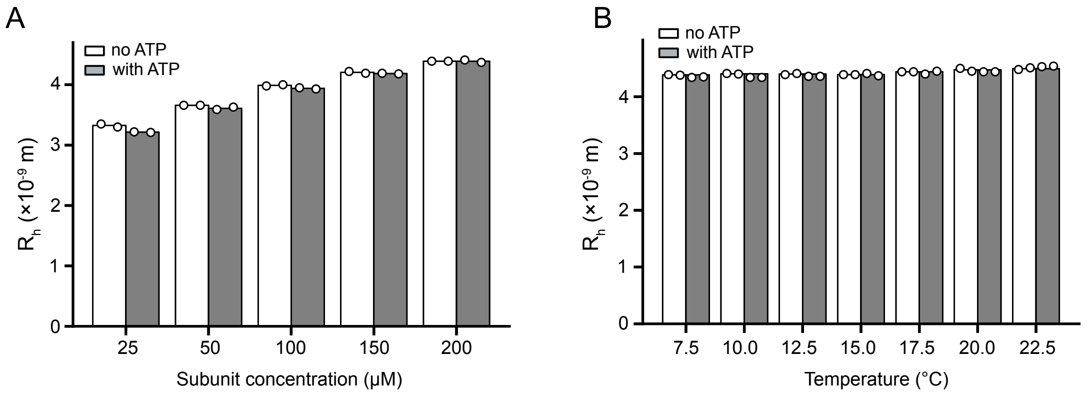


**Fig. S1**. Hydrodynamic radii of $^{\Delta N}$YME1L obtained by dynamic light scattering. (A) Data measured at 15 °C, in the presence and absence of 10 mM ATP, and at various subunit concentrations. (B) Data measured at the subunit concentration of 200 μM, in the presence and absence of 10 mM ATP, and at various temperatures.

25 50 100 150 200

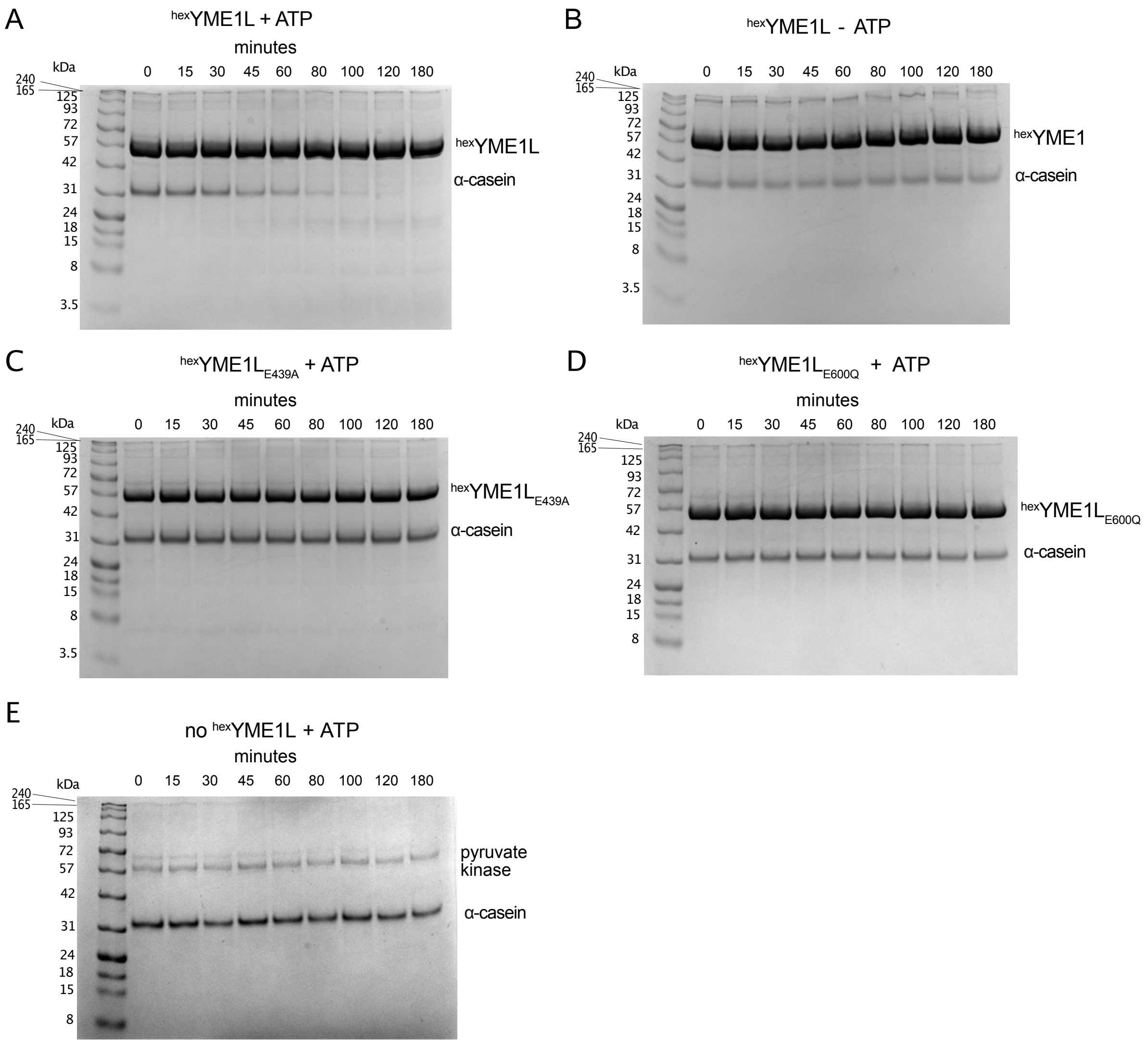


**Fig. S2**. (A-B) Degradation of α-casein by $^{hex}$YME1L in the presence (A) and absence (B) of ATP. (C-D) Degradation of α-casein by $^{hex}$YME1L$_{E439A}$ (Walker-B mutant) (C) and $^{hex}$YME1L$_{E600Q}$ (protease-inhibiting mutant) (D) in the presence of ATP. (E) Degradation of α-casein in the absence of $^{hex}$YME1L as a negative control.

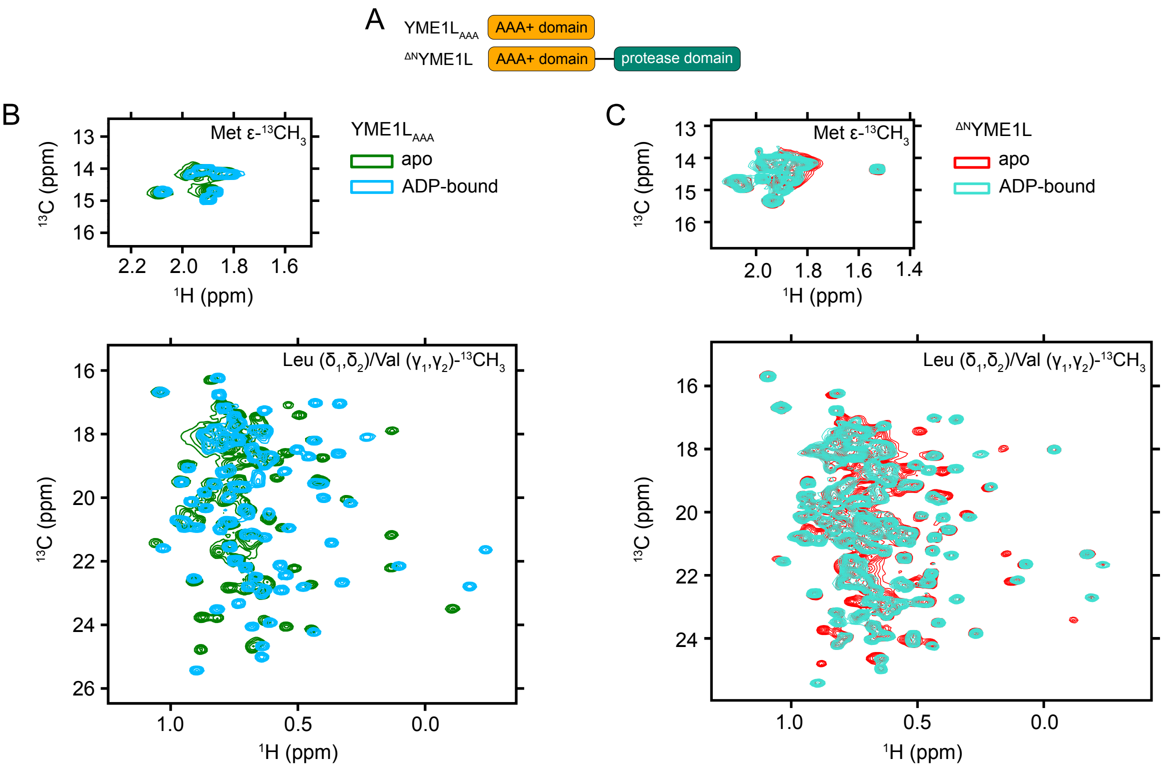


**Fig. S3.** $^{1}$H-$^{13}$C HMQC spectra of YME1L$_{AAA}$ and $^{\Delta N}$YME1L. (A) Domain organization of the two constructs, YME1L$_{AAA}$ and $^{\Delta N}$YME1L, presented in the figure. (B) Overlay of $^{13}$C-$^{1}$H HMQC spectra of [U-$^{2}$H, ILVM-$^{13}CH_3$] labeled YME1L$_{AAA}$ in the presence (blue) and absence (green) of 5 mM ADP. (C) Overlay of $^{13}$C-$^{1}$H HMQC spectra of [U-$^{2}$H, ILVM-$^{13}CH_3$] labeled $^{\Delta N}$YME1L in the presence (green) and absence (red) of 5 mM ADP. Spectra were acquired at 14.1 T and 20 °C.

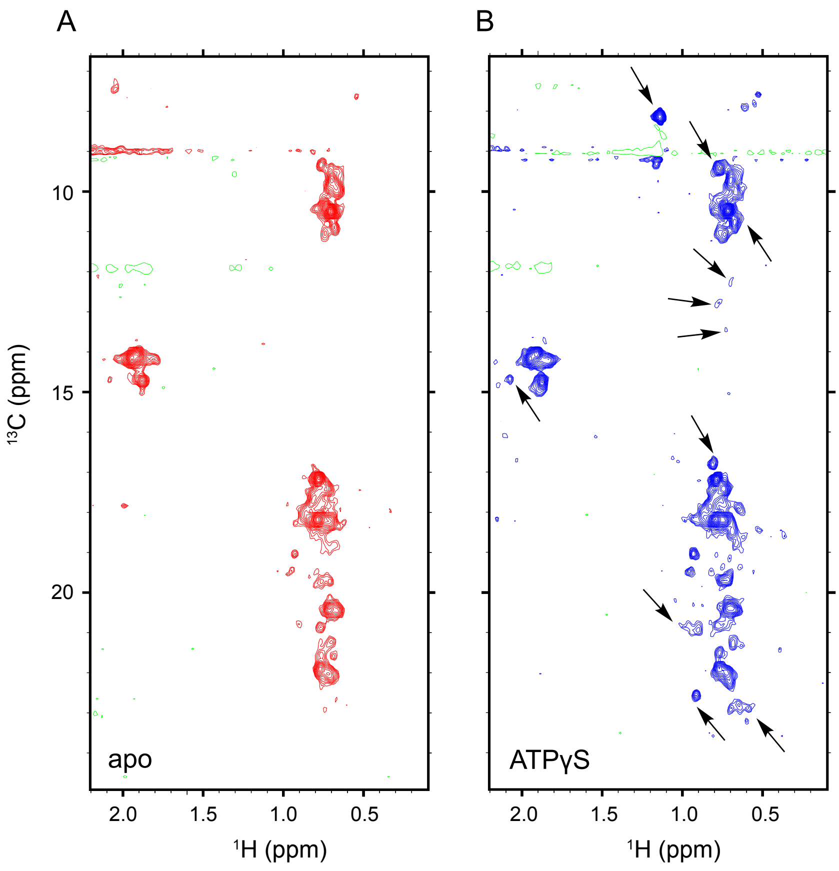


**Fig. S4.** $^{1}$H-$^{13}$C HMQC spectra of [U-$^{2}$H, ILVM-$^{13}CH_3$]-labeled $^{hex}$YME1L$_{AAA}$ in the apo state (A) and in the presence of 20 mM ATPγS (B). Spectra were acquired at 14.1 T and 20 °C. The arrows denote the appearance of new peaks or increase in peak intensity upon the addition of ATPγS .

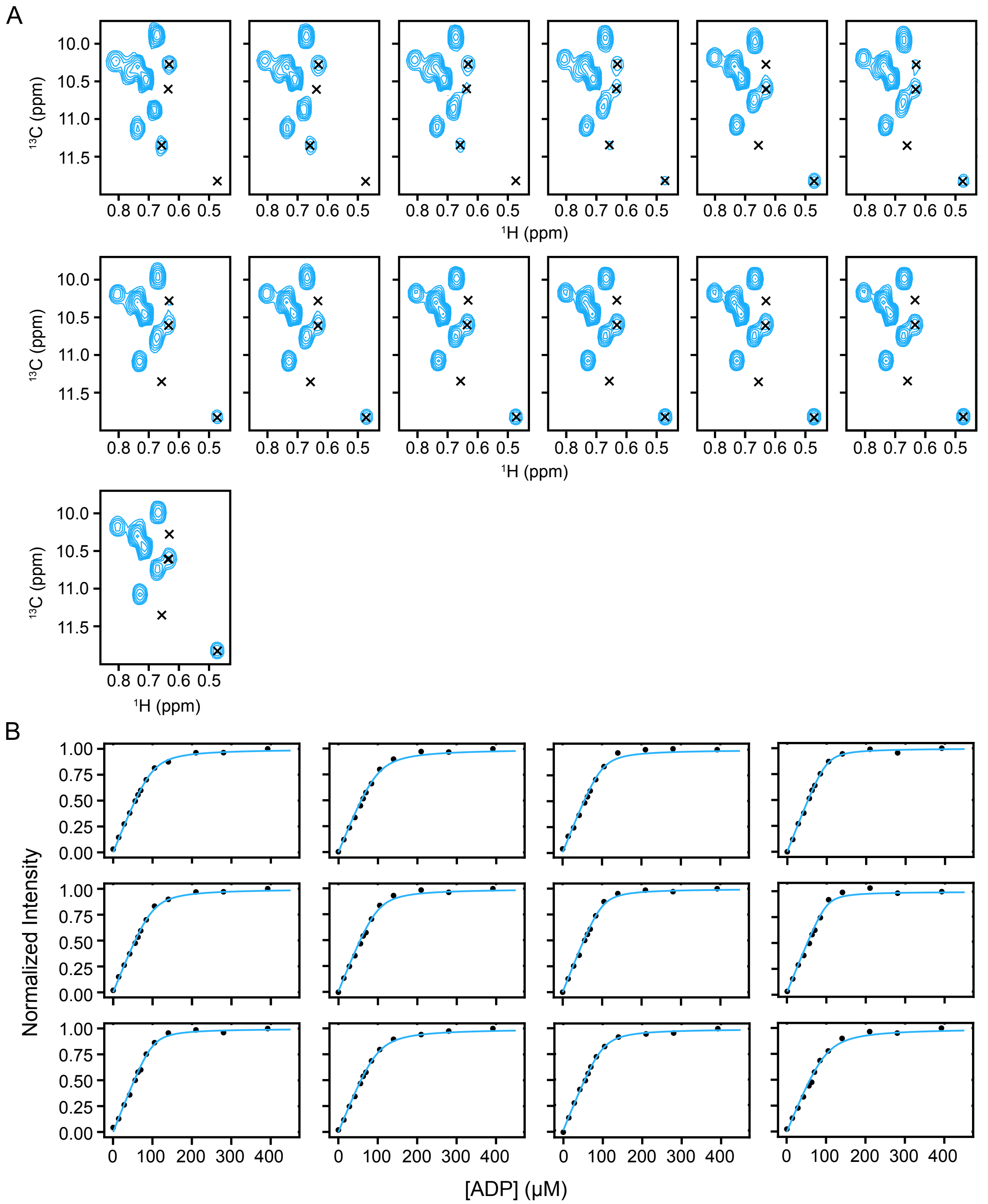


**Fig. S5.** ADP titration into YME1L$_{AAA}$. (A) A selected region of $^{13}$C-$^{1}$H HMQC spectra of [U-$^{2}$H, ILVM-$^{13}$CH$_3$] labeled YME1L$_{AAA}$ in the presence of 0, 14, 28, 42, 56, 63, 70, 84, 105, 140, 210, 280, 392 μM ADP. (B) Titration profiles of selected peaks in the $^{13}$C-$^{1}$H HMQC spectra with increasing concentration of ADP, fitted to a one-to-one binding model for slow chemical exchange. An average $K_d$ value of 4.6 ± 2.3 μM was obtained from the 12 titration profiles in (B) and the 4 peaks in Fig. 2C. Spectra were acquired at 14.1 T and 20 °C.

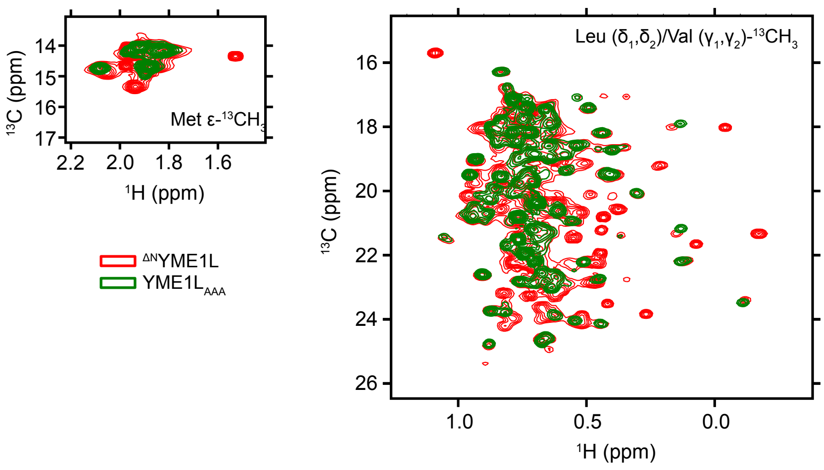


**Fig. S6.** Overlay of $^{1}$H-$^{13}$C HMQC spectra of [U-$^{2}$H, ILVM-$^{13}$CH$_3$] labeled YME1L$_{AAA}$ (green) and $^{\Delta N}$YME1L (red). Spectra were acquired at 14.1 T and 20 °C.

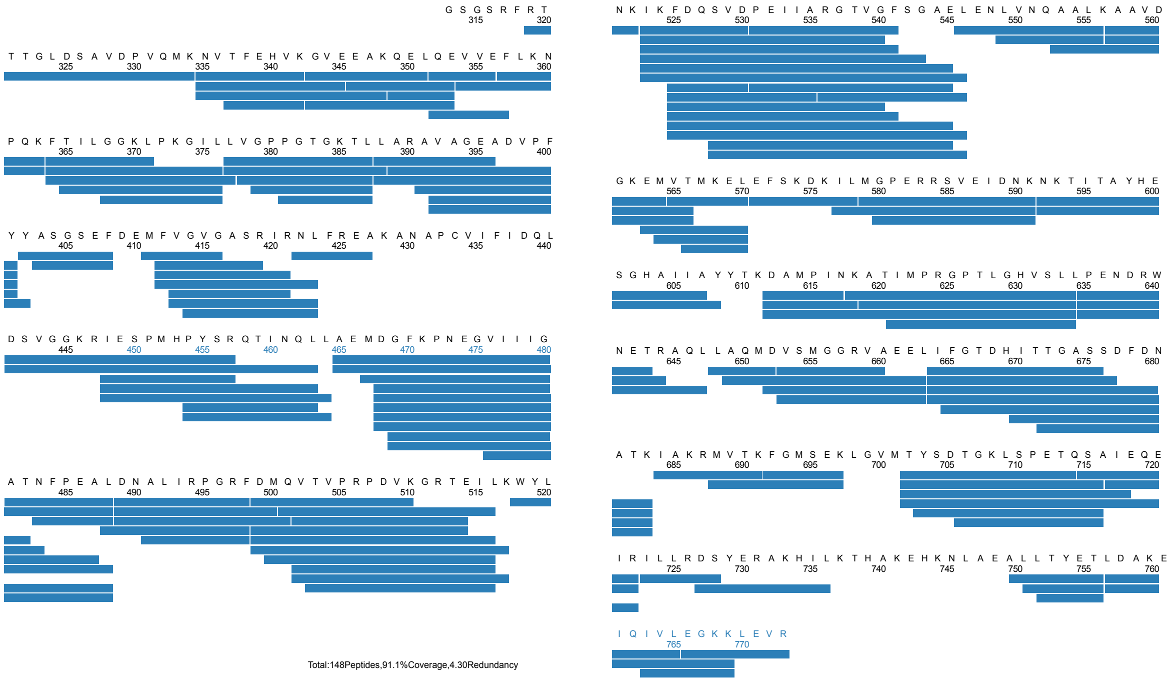


**Fig. S7.** The HDX sequence coverage map of YME1L digested online using a Nepenthesin II column. The sequence coverage is 91% and the sequence redundancy level is 4.3.

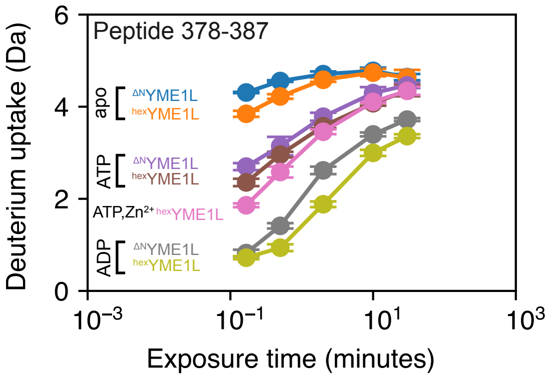


**Fig. S8**. HDX kinetics of the peptide 378-387 for $^{hex}$YME1L and $^{\Delta N}$YME1L in the apo, ATP-bound, and ADP-bound states, as well as the ATP- and $Zn^{2+}$-bound states for $^{hex}$YME1L.

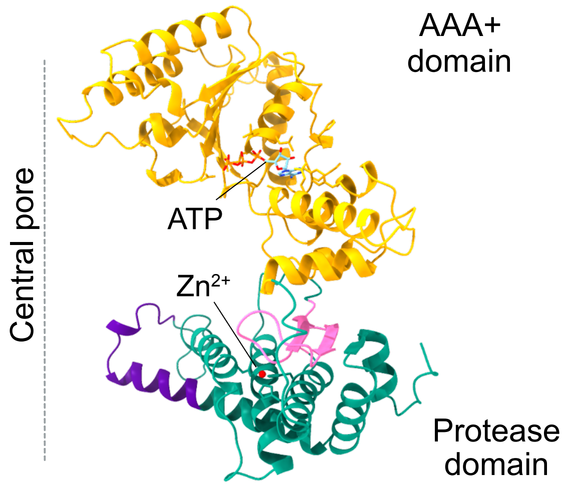


**Fig. S9.** Cartoon representation of the catalytic domain of a YME1L subunit (AlphaFold3) highlighting the location of residues 612-634 (pink) at the domain interface and residues 702-722 (purple) located at the subunit interface facing the central pore.

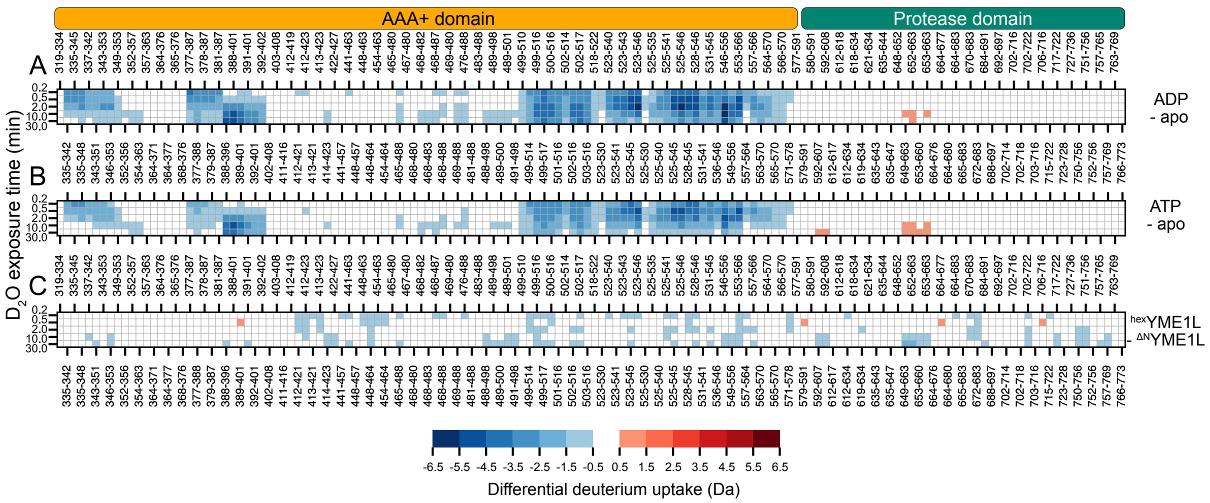


**Fig. S10.** Heatmaps display the differential deuterium uptake of $^{\Delta N}$YME1L bound to (A) ADP and (B) ATP relative to apo $^{\Delta N}$YME1L. In both comparisons, a reduction of deuterium uptake is observed on the nucleotide binding site of the AAA+ domain. (C) The differential deuterium uptake of ATP-bound $^{hex}$YME1L relative to ATP-bound $^{\Delta N}$YME1L shows the additional stabilization given by the hex-tag of $^{hex}$YME1L.

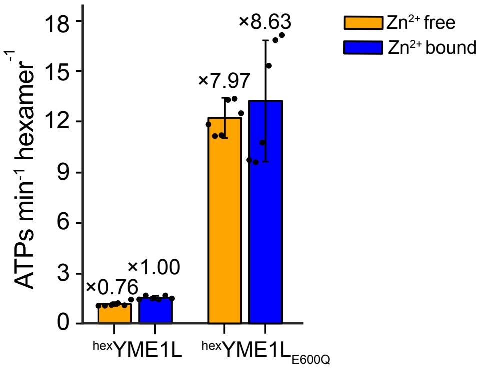


**Fig. S11.** Rates of ATP hydrolysis of $^{hex}$YME1L and $^{hex}$YME1L$_{E600Q}$ in the presence (blue) and absence (yellow) of bound $Zn^{2+}$.

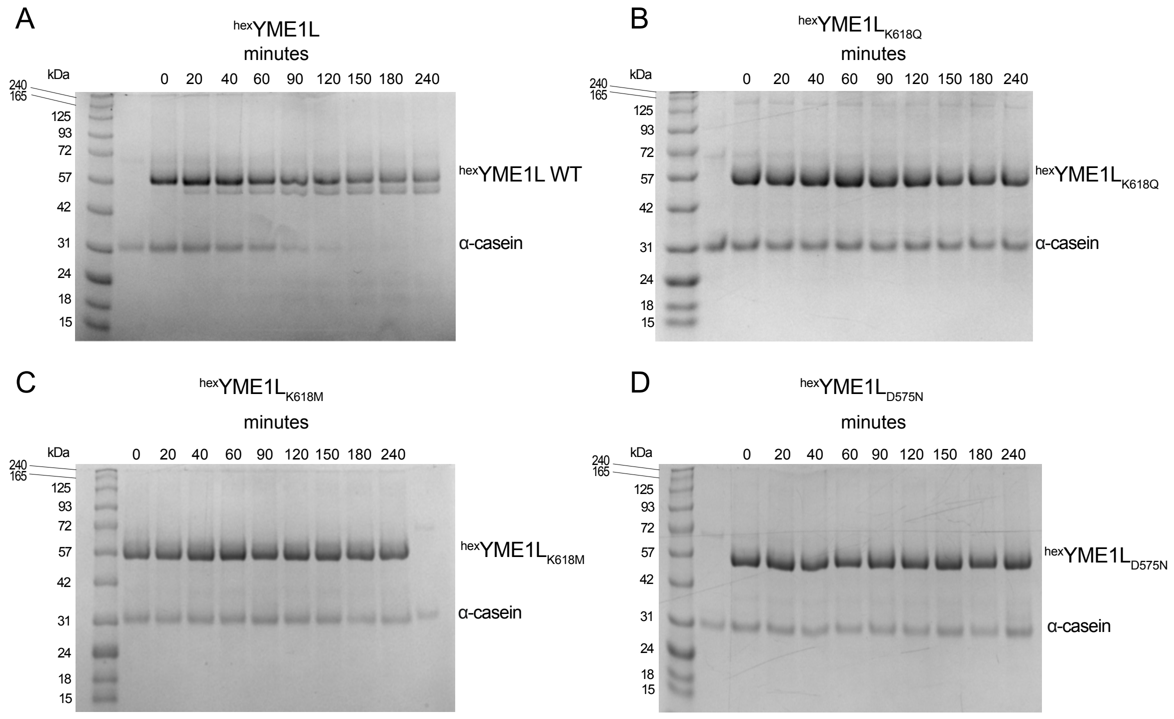


**Fig. S12.** Degradation of α-casein by $^{hex}$YME1L (A), $^{hex}$YME1L$_{K618Q}$ (B), $^{hex}$YME1L$_{K618M}$ (C) and $^{hex}$YME1L$_{D575N}$ in the presence of ATP.

minutes

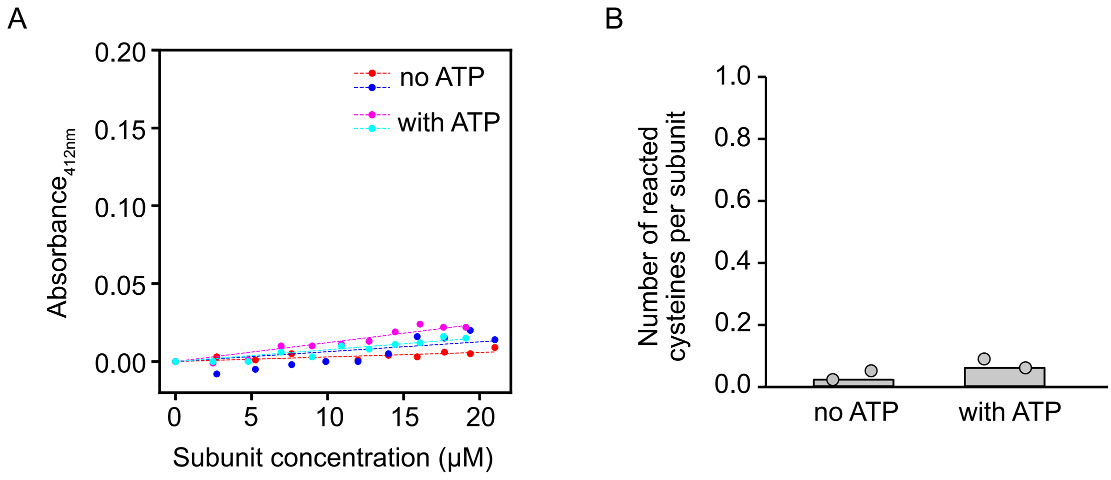


**Fig. S13.** DTNB assay on $^{hex}$YME1L. (A) Absorbance changes at 412 nm with an increasing concentration of $^{hex}$YME1L titrated into the DTNB solution in the presence and absence of 5 mM ATP. (B) The number of reacted cysteine per subunit in the presence and absence of ATP, calculated by fitting the curves in (A) with a linear correlation y = ax.

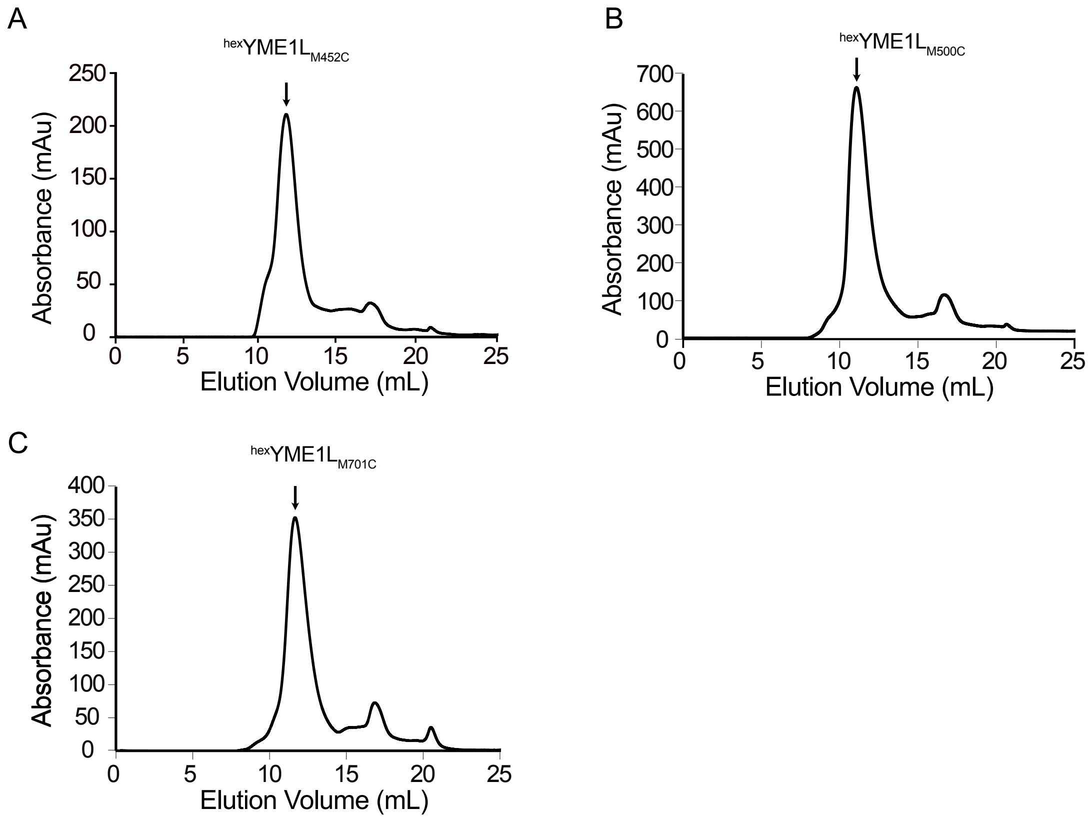


**Fig. S14.** SEC elution profiles of $^{hex}$YME1L$_{M452C}$ (A), $^{hex}$YME1L$_{M500C}$ (B), and $^{hex}$YME1L$_{M701C}$ (C). The elution volumes of the samples are consistent with the size of the hexametric YME1L.

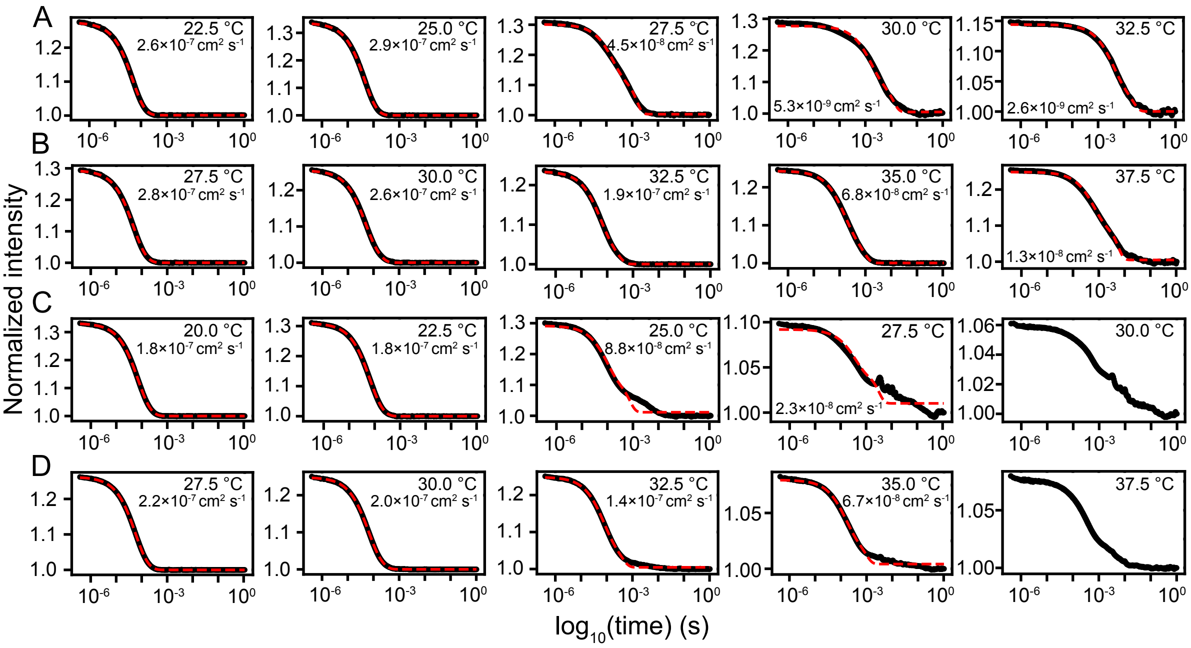


**Fig. S15.** DLS autocorrelation profiles of $^{hex}$YME1L around the critical aggregation temperatures. (A) and (B) show the autocorrelation profiles recorded at the subunit concentration of 25 μM in the absence (A) and presence (B) of 10 mM ATP. (C) and (D) show the autocorrelation profiles recorded at the subunit concentration of 200 μM in the absence (C) and presence (D) of 10 mM ATP. The raw data (black lines) are fit under the assumption of mono-modal distribution of decay rate, with the fits shown by the red dashed lines and the fitted values of diffusion coefficient shown in the graphs. The critical aggregation temperatures were determined to be 25.0, 30.0, 22.5, and 30.0 °C for (A-D), respectively, above which the start of aggregation is indicated by the significant decrease of diffusion coefficient and/or clear deviation from the fits.

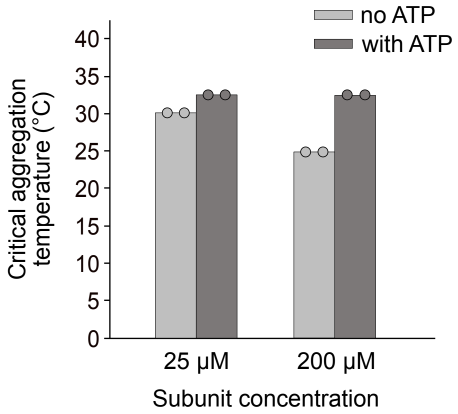


**Fig. S16.** Critical aggregation temperature of $^{\Delta N}$YME1L at the subunit concentrations of 25 µM and 200 µM in the presence and absence of 10 mM ATP, determined by dynamic light scattering.

### Table S1. HDX summary.

| | Apo $^{hex}$YME1L | Apo $^{\Delta N}$YME1L | Apo $^{hex}$YME1L$_{E600Q}$ | ATP-bound $^{hex}$YME1L | ATP-bound $^{\Delta N}$YME1L | ATP-bound $^{hex}$YME1L$_{E600Q}$ | ATP, $Zn^{2+}$-bound $^{hex}$YME1L | ADP-bound $^{hex}$YME1L | ADP-bound $^{\Delta N}$YME1L |
|---|---|---|---|---|---|---|---|---|---|
| HDX reaction details | Final $D_2O$ concentration (v/v): 90%<br>$pH_{corr}$ 7.5, RT | | | Final $D_2O$ concentration (v/v): 90%<br>$pH_{corr}$ 7.5, RT<br>10 mM ATP | | | Final $D_2O$ concentration (v/v): 90%<br>$pH_{corr}$ 7.5, RT<br>10 mM ATP,<br>1 mM $ZnCl_2$ | Final $D_2O$ concentration (v/v): 90%<br>$pH_{corr}$ 7.5, RT<br>10 mM ADP | |
| HDX time course | 0.167, 0.5, 2, 10, 30 min | | | | | | | | |
| Undeuterated controls | 3 | | | | | | | | |
| Back-exchange | 33 % (estimated from Turner, M. *et al* 2024 (*ref*)) | | | | | | | | |
| Number of peptides | 148 | | | | | | | | |
| Sequence coverage | 91 % | | | | | | | | |
| Average peptide length / redundancy | 12.2 ± 4 / 4.3 | | | | | | | | |
| Replicates | 3 technical replicates | | | | | | | | |
| Repeatability | 0.1 Da | | | | | | | | |
| Significant differences | 0.5 Da | | | | | | | | |
